# GRASPS: a simple-to-operate translatome technology reveals omics-hidden disease-associated pathways in TDP-43-related amyotrophic lateral sclerosis

**DOI:** 10.1101/2024.03.04.583294

**Authors:** Ya-Hui Lin, Jennifer E. Dodd, Luisa Cutillo, Lydia M. Castelli, Simeon R. Mihaylov, Karl Norris, Adrian Higginbottom, Matthew J. Walsh, Johnathan Cooper-Knock, J. Robin Highley, Ilaria Granata, Caroline A. Evans, Mario R. Guarracino, Susan G. Campbell, Mark J. Dickman, Pamela J. Shaw, Marta Milo, Guillaume M. Hautbergue

## Abstract

Transcriptomes and translatomes measure genome-wide levels of total and ribosome-associated RNAs. A few hundred translatomes were reported over >250,000 transcriptomes highlighting the challenges of identifying translating RNAs. Here, we used a human isogenic inducible model of TDP-43-linked amyotrophic lateral sclerosis, which exhibits altered expression of thousands of transcripts, as a paradigm for the direct comparison of whole-cell, cytoplasmic and translating RNAs, showing broad uncoupling and poor correlation between disease-altered transcripts. Moreover, based on precipitation of endogenous ribosomes, we developed GRASPS (Genome-wide RNA Analysis of Stalled Protein Synthesis), a simple-to-operate translatome technology. Remarkably, GRASPS identified three times more differentially-expressed transcripts with higher fold changes and statistical significance, providing unprecedented opportunities for data modeling at stringent filtering and discovery of previously omics-missed disease-relevant pathways, which functionally map on dense gene regulatory networks of protein-protein interactions. Based on its simplicity and robustness, GRASPS is widely applicable across disciplines in the biotechnologies and biomedical sciences.

## Introduction

Transcriptomics has been widely used to interrogate prokaryotic and eukaryotic transcriptomes over the past 25 years. However, up-regulation of mRNA levels does not necessarily correlate with increased levels of the corresponding proteins, but rather down regulation as cells attempt to compensate for the down-regulated proteins. Genome-wide investigation of human gene expression involves quantifying the expression levels of ∼200,000 transcripts assigned to ∼58,000 genes, including 145,571 protein-coding transcripts expressed from 19,591 protein-coding genes^1–3^ using next generation RNA sequencing technologies (RNA-seq). Approximately 70,000 proteins are annotated in *Ensembl*^4^ while post-translational modifications, which modulate the function or activity of proteins, give rise to hundreds of thousands of protein variants^5^. On the other hand, the abundance of mammalian proteins is mostly regulated by their biosynthesis and turnover/degradation^6, 7^ while less than one third can be attributed to total mRNA concentrations^8^. The genome-wide identification of mRNA molecules associated with ribosomes in translatomes is thus expected to functionally reflect the directionality of protein expression changes in developing tissues, cellular homeostasis or diseases. Methodologies identifying translatomes include polysome profiling, ribosome profiling and ribosome affinity purification^9^.

Polysome profiling, the first methodology developed 20 years ago to investigate the translatome of the yeast *Saccharomyces cerevisiae*, is based on the qualitative separation of polysomes through sedimentation in a sucrose gradient^10^. Extracted RNA molecules are then identified using qRT-PCR, micro-arrays or RNA-seq. More recently, ribosome profiling allowed genome-wide mapping of ribosomes onto RNA molecules at near nucleotide resolution through sequence identification of ribosome-protected RNA fragments generated upon limited RNAse treatment^11, 12^. This technique greatly informed our knowledge of the cellular translational activity, including association of ribosomes with short open reading frames in long intergenic non-coding (linc) RNA^13^, annotating coding parts of genomes^12^ and the finding that miRNAs mainly trigger decreased mRNA levels rather than translational inhibition in mammals^14^. On the other hand, translating ribosome affinity purification (TRAP) allows cell type-specific purification of tagged ribosome subunits^15^. It was particularly successfully applied *in vivo* to identify translatomes specific for various neuronal cell populations through promoter-restricted expression of GFP-, YFP- or HA-tagged ribosomal proteins L10a or L22 in rodent brains^16–18^.

However, current translatome technologies present technical challenges as illustrated by the relatively low number of publications compared to transcriptome studies (∼500 versus >250,000 over the past 20-25 years). Major drawbacks include^9, 19^: (i) low throughput; (ii) large volume and high variability in sucrose gradient-dependent size fractionation of polysomes; (iii) engineering of transgenic models overexpressing tagged ribosome subunits which likely alter functionality and does not allow purification of the cellular fraction of untagged ribosomes; (iv) requirement for “freezing” inhibitors, such as cycloheximide, to maintain ribosome-mRNA associations during the course of purification but which cause translational stress artefacts^20^. It is generally assumed that RNAs associated with ribosomes are translated into proteins. However, non-translated RNA molecules can be co-purified from paused/stalled ribosomes or by indirect binding to ribosomes via other RNA-processing proteins. For example, the multi-functional DNA/RNA-binding protein TDP-43 (TAR DNA-binding protein-43) interacts with ribosomes under stress^21, 22^, potentially contaminating ribosome preparations with hundreds of indirectly bound mRNA molecules transported by TDP-43^23, 24^. Non-translating, initiating and elongating ribosomes are all purified from whole-cell extracts used to perform ribosome affinity purification or ribosome profiling. In the latter case, ribosome-protected RNA fragments are also particularly short (approximately 25-30 nucleotides) presenting added challenges for mapping reads to repeated sequences and alternatively spliced transcripts^19^.

Despite the aforementioned limitations, translatomes present unique opportunities to interrogate the expression of genes at a functional level, benefiting from the high genome coverage permitted by RNA-seq (identifying 60-80% RNAs) while the dynamic molecular range of proteomes is limited by protein abundance and the lower number of proteins quantified by mass spectrometry^25^ (typically ∼10% expressed proteins). The Human Protein Atlas project, a world-wide initiative started 20 years ago for the characterization of all human proteins, showed a low correlation between transcriptomes and proteomes in 29 human tissues in which the expression of genes was characterized through the identification of 18,072 transcripts and 13,640 proteins^25^ due to: (i) the aforementioned difference in the dynamic molecular ranges between RNA-seq and mass spectrometry leading to limited detection of lower-expressed proteins; (ii) the number of molecules of proteins produced per molecule of mRNA depends on the abundance of the given mRNA i.e. higher for mRNAs with increased expression^25^. A few studies also reported a low correlation between transcriptomes and translatomes, showing extensive uncoupling across 20 paired mammalian transcriptome-translatome datasets^26^. More recently, gene ontology indicated that the network of genes built at transcriptional level is rewired at translational level^27^ while total mRNA levels are less dynamic than translating mRNAs^28^. Accordingly, the translatome is better correlated with the proteome in a parasite study^29^.

Here, we developed a novel translatome methodology easily scalable to any biochemistry or molecular biology laboratory and we provide 4 high-depth RNA-seq datasets as a resource for the direct comparison of total, cytoplasmic and translating RNAs using transcriptome and translatome approaches. For this investigation, we engineered human cells with isogenic inducible expression of TDP-43 Q331K, a DNA/RNA-binding mutant protein causing the neurodegenerative disease amyotrophic lateral sclerosis (ALS)^30^ and which causes altered expression of hundreds to thousands of RNAs in mouse and human ALS brains^31–35^. Despite a large number of transcriptome studies, the functional consequences of widespread dysregulation of RNA metabolism in ALS remain poorly investigated.

## Results

### Engineering a human isogenic TDP-43 ALS-inducible cell model

We engineered a plasmid expressing the TDP-43 Q331K ALS mutant^30^ under a tetracycline-regulated cytomegalovirus (CMV) promoter prior to integration into a human embryonic kidney (HEK) cell line harboring the Flp-In™ T-REx system which allows the generation of stable cell lines by homologous recombination at the engineered Flp Recognition Target (FRT) locus (**Methods**). The backbone plasmid was also integrated into the HEK Flp-In™ cells to generate an isogenic sham control line^36–40^. Adding tetracycline to the cell culture for 48 hours leads to specific promoter de-repression and induction of the TDP-43 Q331K mutant protein detected alongside the endogenous TDP-43 protein (**Fig. 1a**). Quantification showed moderate overexpression with a 3-fold increase in total TDP-43 proteins and ALS-relevant co-expression of both wild type and mutant proteins (**Fig. 1b**). The transcriptional induction of TDP-43 Q331K was further validated in tetracycline-treated lines at mRNA level (**Fig. 1c**). Immunofluorescence microscopy further confirmed the low level of TDP-43 overexpression upon induction, while endogenous TDP-43 and TDP-43 Q331K proteins remain predominantly nuclear (**Fig. 1d**), in agreement with a previous study which reported a mass spectrometry investigation of TDP-43 complexes in isogenic HeLa Flp-In™ cells^41^. The growth of non-induced and tetracycline-induced sham control and TDP-43 Q331K cell lines was measured over 15 days, showing that expression of TDP-43 Q331K specifically inhibits the cell proliferation of this model of TDP-43-mediated cytotoxicity (**Fig. 1e**).

**Fig. 1.**
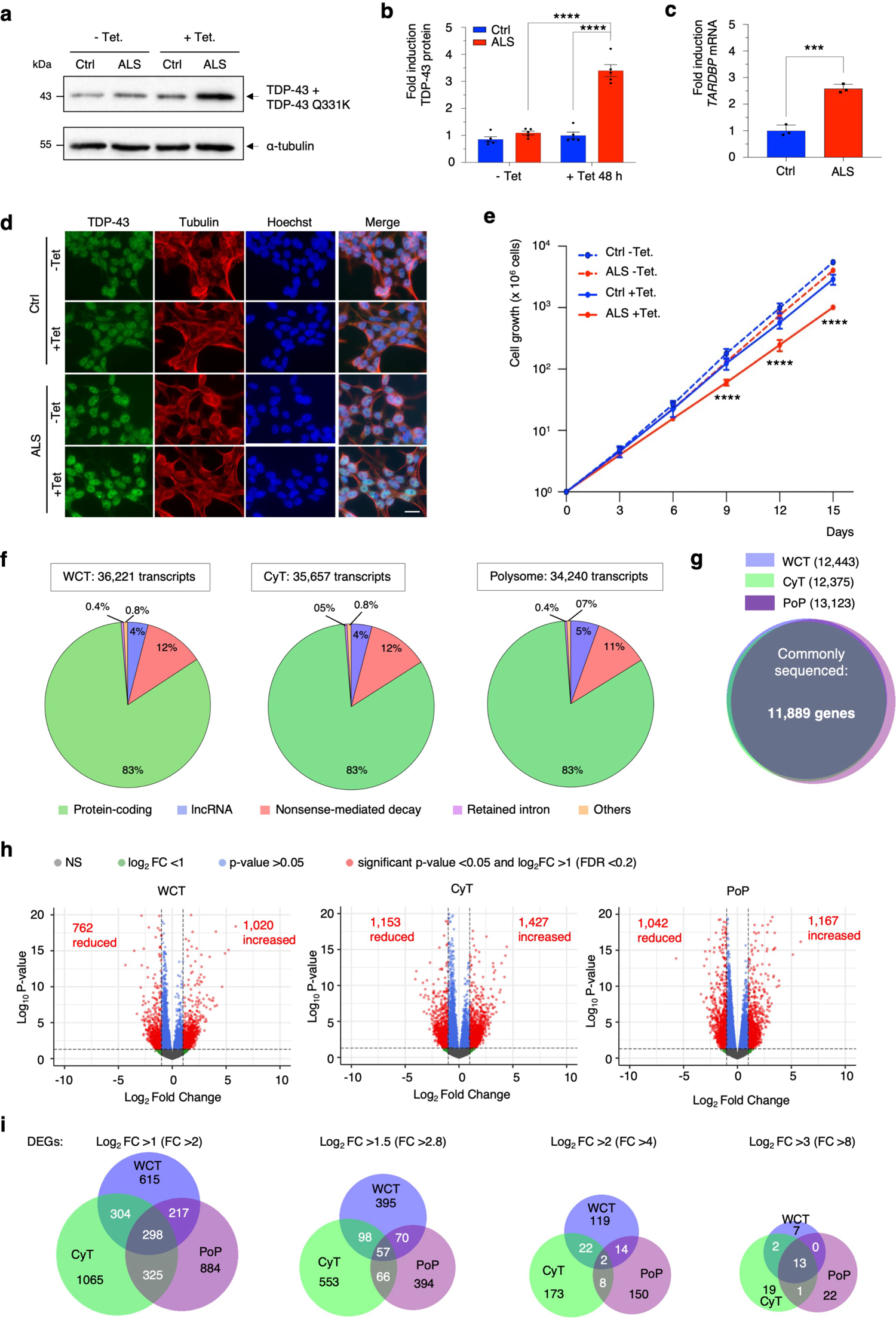
Genome-wide RNA expression and translation in human isogenic TDP-43 ALS-inducible cells. Control (Ctrl) and TDP-43 Q331K (ALS) cell lines were treated with tetracycline to induce transgene expression for 48 hours. **(a)** TDP-43 and α-tubulin protein levels were analyzed by western immunoblotting with (+Tet.) or without (-Tet.) tetracycline induction. **(b)** Quantification of TDP-43 wild type and mutant protein expression following normalization to α-tubulin in three biological replicates (Mean ± SEM; two-way ANOVA with Tukey’s correction for multiple comparisons; ****: p<0.0001; N=3). **(c)** qRT-PCR quantification of *TARDBP* mRNA encoding TDP-43 proteins following normalization to *U1 snRNA* in three biological replicates (Mean ± SEM; unpaired t test; ***: p=0.0005; N=3). **(d)** Immunofluorescence microscopy. Cells were stained with Hoechst to label nuclei (blue), TDP-43 (green) and α-tubulin (red). Scale bar: 20 μm. **(e)** Growth curves. Non-induced (-Tet.) and tetracycline-induced (+Tet.) cells were split every 72 hours for the indicated number of days. Total cell numbers were counted before splitting and quantified in three biological replicates (Mean ± SEM; two-way ANOVA with Tukey’s correction for multiple comparisons; ****: p<0.0001; N=3). **(f)** Pie charts representing the distribution and main categories of annotated transcripts sequenced in WCT, CyT and PoP. **(g)** Venn diagram of sequenced genes in WCT, CyT and PoP. **(h)** Volcano plots representing differentially-expressed transcripts according to p-values and fold changes (FC) in WCT, CyT and PoP. NS: non-significant. Red labels indicate the numbers of significantly down- and up-regulated transcripts. **(i)** Venn diagrams representing the distribution of DEGs in WCT (blue), CyT (green) and PoP (purple) at increasing FC thresholds.

### Widespread and uncoupled alteration of total, cytoplasmic and translating RNAs in human TDP-43 ALS-inducible cells

Sham control (Ctrl) and TDP-43 Q331K (ALS) cell lines were induced with tetracycline for 48 hours prior to cellular fractionation and purification of whole-cell, cytoplasmic and polysome-associated RNAs. Total RNAs were extracted from whole lysed cells while the cytoplasmic fraction was isolated upon hypotonic lysis^40, 42, 43^. The quality of the fractionation was validated by the absence of detectable nuclear contamination in the cytoplasmic fractions probed with the chromatin remodeling factor SSRP1 (**Supplementary Fig. 1a**). We also performed polysome profiling experiments to isolate the actively translating fraction of RNAs (**Supplementary Fig. 1b**). RNA samples were used for the preparation of RNA sequencing libraries which were subjected to high depth next generation *Illumina* sequencing (**Methods**) covering the human transcriptome and translatome by 40-fold and 400-fold respectively – 10% of the genome being transcribed^44, 45^ while less than 2% is considered translated^46^ (**Supplementary Data 1**). Over 34,000 annotated transcripts were sequenced in the whole cell transcriptome (WCT), cytoplasmic transcriptome (CyT) and polysome profiling (PoP) experiments (**Supplementary Data 2 tabs 1-3**). Protein-coding mRNAs (83%), nonsense-mediated decay (NMD) transcripts (11-12%) and long non-coding (lnc) lncRNAs (4-5%) were predominantly sequenced with the same proportion in all datasets, providing further evidence that many lncRNAs are translated or regulate translation^47, 48^ while in agreement with these data, NMD is linked to translation termination^49^ (**Fig. 1f**). These transcript isoforms map 11,889 genes which were commonly sequenced across all conditions, thus validating the identification of high-quality datasets without notable sequencing bias (**Fig. 1g**). Sequencing reads were aligned on GRCh38 (hg38) using Bowtie^50^. Changes in RNA expression levels were quantified between the transcriptomes and translatomes of the tetracycline-induced Ctrl and ALS cell lines using BitSeq^51^ and Limma^52^. For the statistical analysis, differentially-expressed (DE) transcripts were filtered for fold change (FC) log2FC>1 (equivalent to FC>2), p-value p<0.05 and false discovery rate FDR<0.2. Lists of differentially-expressed transcripts and genes (DEGs) are provided in **Supplementary Data 3 tabs 1-3** for annotated transcripts in WCT/CyT transcriptomes and PoP translatomes respectively. Expression levels of 2,200–2,750 total, cytoplasmic and translating RNAs are altered upon induction of TDP-43 Q331K in the human ALS disease model with 900–1,500 transcripts being either up- or down-regulated in each dataset (**Fig. 1h**). Overall, the expression of 1434, 2101 and 1724 genes is altered at total, cytoplasmic and translating levels (**Supplementary Data 3**, columns labeled DEGs) with only 20% or 30% of gene expression changes shared across all or with another dataset respectively, in agreement with recent reports^26–28^ (**Fig. 1i**). However, this study also highlights a poor correlation between cytoplasmic transcriptomes and translatomes which are counter-intuitively distant from each other. Increasing the stringency of filtering using higher FC thresholds leads to drastic reduction of significantly altered genes in all datasets with the identification of only 150-200 DEGs at FC>4 and 20 or less at FC>8 (**Fig. 1i**), a known bottleneck of RNA-seq studies typically requiring filtering data on low fold change thresholds of 1.3-2.0 which may not reflect significant biological effects.

### GRASPS: a translatome technology based on stringent pelleting of ribosomes

Despite higher statistical power for the above polysome profiling with a genome coverage of 400 fold compared to 40 fold for the transcriptomes (less than 2% of the genome considered coding^46^ over 10% transcribed^44, 45^; **Supplementary Data 1**), the number of DEGs did not noticeably increase for the PoP translatome (**Fig. 1h**, **Supplementary data 3 tabs 1-3**). Other translatome studies related to polysome^26^, ribosome^53^ or TRAP^54^ profiling experiments identified a similar range of expression changes in varied models. We hypothesized that higher experimental variability in longer and more complex to operate translatome technologies is a potential factor for reduced statistical power and identification of translating RNA changes. Therefore, to avoid using a translation inhibitor and sucrose gradients or immunoprecipitation, we developed an alternative approach based on ultraviolet (UV) irradiation and stringent pelleting of ribosomes covalently crosslinked to RNAs. Briefly, this protocol involves hypotonic lysis of cells to isolate a cytoplasmic fraction which is subjected to UV irradiation and differential centrifugation in high salt to remove mitochondria and further pelleting of ribosome:RNA complexes through a sucrose cushion. This protocol is simple and time efficient with ribosome-associated RNA being purified in one tube within 6 hours, in contrast to approximately 24-30 hours and multiple RNA samples per condition while using a sucrose gradient in polysome profiling (**Fig. 2, Methods**). The buffer conditions were optimized for selective enrichment of actively translating RNA molecules using transient incubation of human HEK cells at 42°C, which is well-known to induce rapid transcription and translation of the *HSPA1A* mRNA encoding cytoplasmic-inducible heat shock protein HSP72/HSP70.1^55, 56^. Accordingly, a time course experiment showed that expression levels of the HSP72 protein is markedly increased 2 hours after heat shock, while those of the control GAPDH protein (glyceraldehyde-3-phosphate dehydrogenase) are not changed (**Fig. 3a**). Quantification of western blots and total RNA levels by qRT-PCR indicated that both HSP72 protein and *HSPA1A* mRNA are induced by approximately 5-fold over the 2-hour time course, while control GAPDH levels do not significantly vary (**Fig. 3b-c**). Consistently, GRASPS measured a 5-fold induction in the translation of the *HSPA1A* mRNA while translating *GAPDH* transcripts remained unaltered after the heat shock (**Fig. 3d**). Mass spectrometry analysis of GRASPS-purified complexes identified 120 proteins (**Supplementary Data 4**). Strikingly, 70 and 6 of these were respectively ribosomal proteins and involved in translation. Overall, the 70 ribosomal proteins and eukaryotic elongation factor 2 (eEF2) map over the 82 subunits of the human 80S ribosome structure^57^, highlighting a high degree of purity and stoichiometry (**Figs. 3e-f**). Gel electrophoresis analysis confirmed enrichment in typically low molecular weight ribosomal proteins, while it also validated the mass spectrometry data and stringent purification of ribosomes with absence of detectable accessory proteins such as eukaryotic initiation translation factor eIF4A (**Fig. 3g**).

**Fig. 2.**
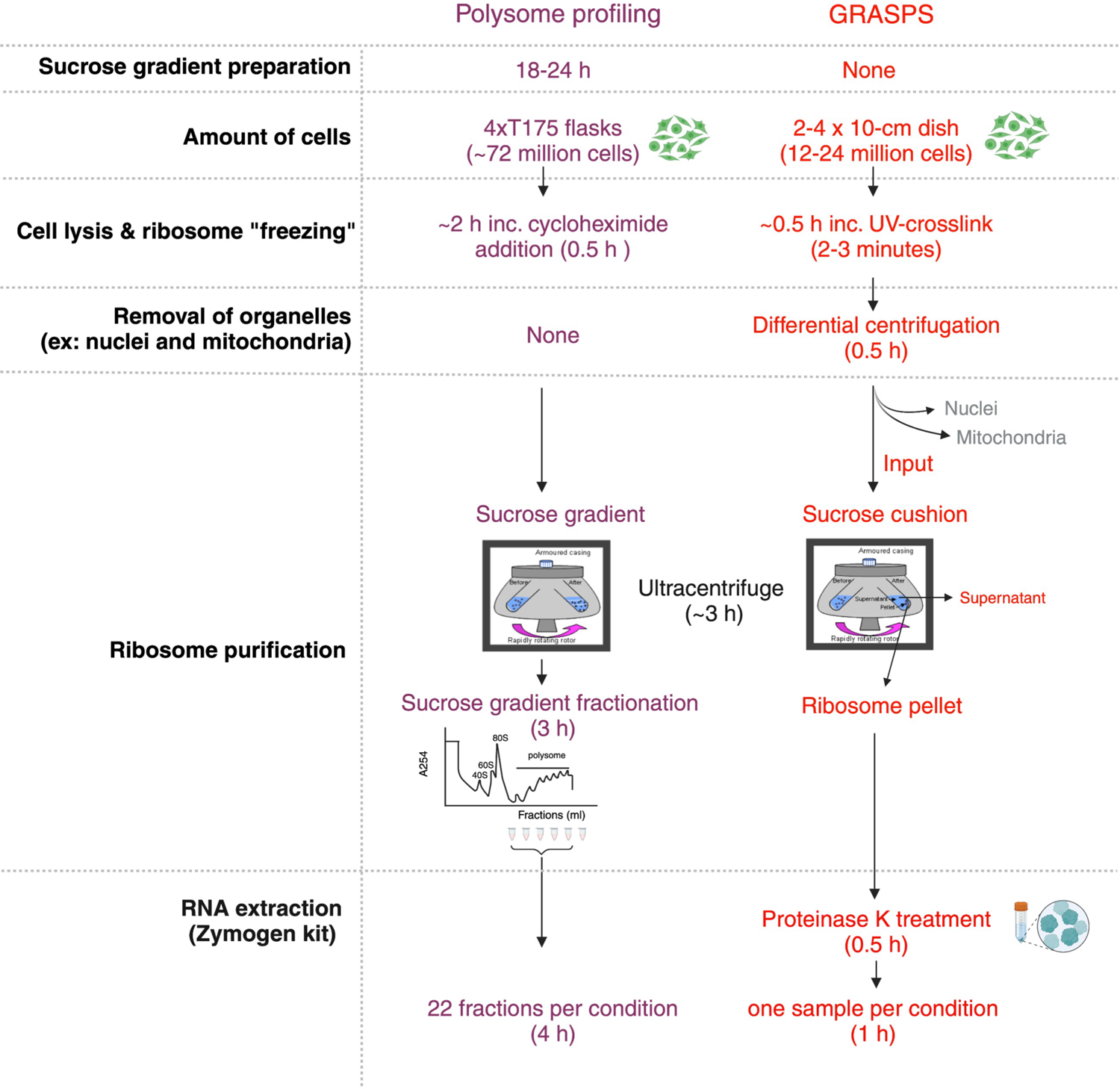
Diagrammatic overview of the GRASPS methodology compared to polysome profiling. UV irradiation is carried out in GRASPS to rapidly stall translating RNA-ribosomes complexes, thus bypassing the reliance on the addition of translation inhibitors such as cycloheximide used in the other translatome technologies. UV cross-linked RNA-ribosomes are further pelleted by ultracentrifugation prior to proteinase K treatment and RNA extraction under 6 hours in contrast to lengthy and delicate sucrose gradient preparation and fractionation which also requires extraction of RNA from multiple fractions for each experimental condition (approximately 24-30 hours).

**Fig. 3.**
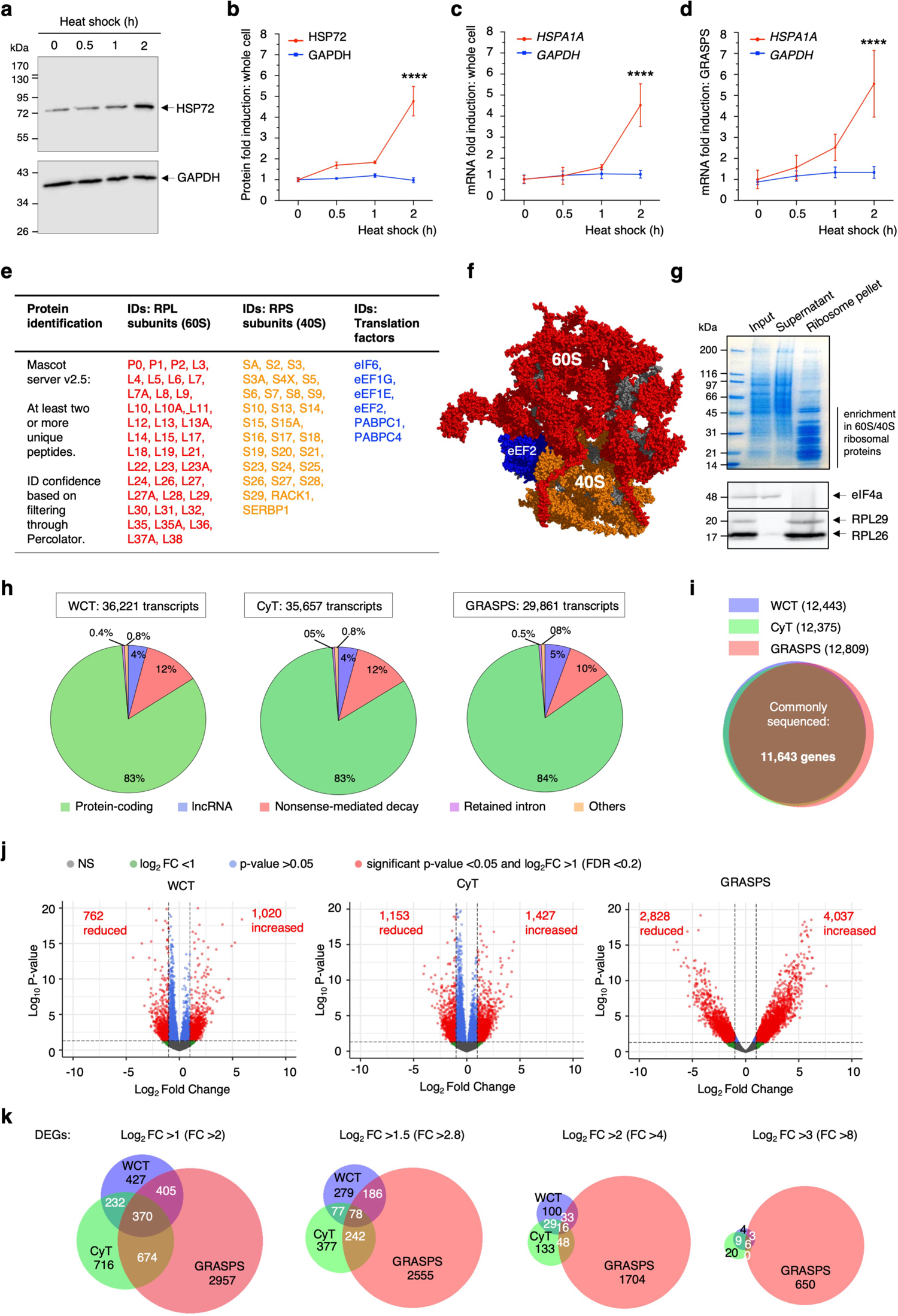
Development of the GRASPS translatome technology and application to the human TDP-43 ALS-inducible cells. **(a)** Heat-shock inducible HSP72 and control GAPDH protein levels were analyzed by western immunoblotting in HEK cells shifted to 42°C for the indicated times. **(b)** Quantification of HSP72 and GAPDH protein levels in four biological replicates (Mean ± SEM; two-way ANOVA with Tukey’s correction for multiple comparisons; ****: p<0.0001; N=4). **(c-d)** qRT-PCR quantification of total (c) and GRASPS-purified (d) *GAPDH* and *HSPA1A* mRNA encoding the HSP72 protein in heat-shocked HEK cells in four biological replicate experiments (Mean ± SEM; two-way ANOVA with Tukey’s correction for multiple comparisons; ****: p<0.0001; N=4). **(e)** Liquid chromatography tandem mass spectrometry (LC-MS/MS) analysis of GRASPS-purified samples identified 115 proteins with a Mascot score >25 and at least two different peptides. Among these, 76 were large (RPL) and small (RPS) ribosomal proteins. 5 additional RPLs were identified manually. **(f)** GRASPS-identified RPL/RPS subunits (70) and eEF2 were mapped onto the cryo-EM structure of the 80S human ribosome^57^ (Protein Data Bank accession 4V6X; 87% coverage, 71 over 82 proteins). Identified large 60S and small 40S ribosome subunits are labeled in red and orange respectively. Elongation factor eEF2 is labeled in blue. 44 other proteins including RNA-binding proteins (hnRNPs) and other abundant cytoskeleton proteins typically contaminating mass spectrometry samples were also identified (Supplementary Data 4). **(g)** SDS-PAGE stained with coomassie blue highlighted enrichment in small molecular weight ribosomal proteins in GRASPS-purified ribosome pellets in comparison to input (total protein extract) and sucrose cushion supernatant. Samples were also analyzed by western immunoblotting probed for eIF4A, RPL26 and RPL29. **(h)** Pie charts representing the distribution and main categories of annotated transcripts sequenced in WCT, CyT and GRASPS. **(i)** Venn diagram of sequenced genes in WCT, CyT and GRASPS. **(j)** Volcano plots representing differentially-expressed transcripts according to p-values and fold changes (FC) in WCT, CyT and GRASPS. NS: non-significant. Red labels indicate the numbers of significantly down- and up-regulated transcripts. **(k)** Venn diagrams representing the distribution of DEGs in WCT (blue), CyT (green) and GRASPS (salmon) at increasing FC thresholds.

### Applying GRASPS to the identification of gene expression changes in the human TDP-43 ALS-inducible cell model

Tetracycline-induced control and TDP-43 Q331K cell lines were subjected to GRASPS and RNA-seq with a translating genome coverage of 180-fold (**Supplementary Data 1**). Approximately 30,000 annotated transcripts were sequenced in GRASPS (**Supplementary Data 2 tab 4**). Compared to previous datasets, the same proportion of protein-coding mRNAs (84%), NMD transcripts (10%) and lncRNAs (5%) were sequenced (**Fig. 3h**). These transcript isoforms map 11,643 genes which were commonly sequenced across all conditions without notable sequencing bias (**Fig. 3i**). Lists of differentially-expressed annotated transcripts and genes, quantified using same thresholds as above, are provided in **Supplementary Data 3 tab 4**. In contrast to the transcriptomes and the PoP translatome, GRASPS identified over 3 times more DEGs at FC>2 with expression levels of 4077 and 2583 transcripts respectively up- and down-regulated upon induction of TDP-43 Q331K in the human ALS disease model (**Fig. 3j**). Strikingly, the vast majority of transcripts are all significantly differentially-expressed (red dots) having very low p-values and high fold changes compared to the transcriptomes or the PoP (**Fig. 1h**) that show a large proportion of non-significantly differentially-expressed transcripts (blue and green dots). Overall, 4406 annotated and 4117 protein-coding genes were differentially expressed in GRASPS. Reminiscent of the PoP translatome integration in **Fig. 1i**, only a small proportion of GRASPS-identified DEGs are shared with the transcriptomes (**Fig. 3k**). However, in contrast to the striking reduction observed while increasing the filtering threshold in the other datasets, the expression of over 650 altered genes is still significantly quantified as dysregulated with FC>8 in GRASPS (**Fig. 3k**), thus allowing the characterization of highly-altered pathways and gene expression changes which are expected to have higher functional relevance at the biological level.

### Integrating GRASPS translatomes with transcriptomes and polysome profiling

Reminiscent of the above RNA-seq investigations, 11,581 genes were commonly sequenced across the 4 datasets (**Supplementary Fig. 2**). Compared with the WCT transcriptome and the Venn diagrams in **Figs. 1i** and **2k**, GRASPS identified approximately 3, 5, 10 and 26 times more gene expression changes when DEGs were respectively filtered for fold changes over 2.0, 2.8, 4.0 and 8.0 while the sizes of the CyT transcriptome and PoP translatome remained similar to WCT ranging between 0.9 - 1.4 (**Fig. 4a**). This indicates that GRASPS is a much more sensitive technology compare to transcriptomics and profiling of actively translated RNAs. The majority of gene expression changes is rather specific for the transcriptomes or translatomes, showing broad uncoupling between total, cytoplasmic and translating RNAs. 30-55% gene expression changes are only quantified in one of the transcriptomes or the PoP translatome while, due to its larger size, 75-90% of DEGs are only identified in GRASPS (**Fig. 4b**). Interestingly, GRASPS and CyT share the highest number of DEGs with the same direction of changes (20.8%) while only 2.9% with the WCT, in expectation with the cytoplasmic pool of RNAs reflecting better the translated fraction. Only 113 genes over the dysregulation of thousands are commonly affected in all datasets (**Supplementary** Fig. 3**, Supplementary Data 5 tab 1**). To assess the degree of correlation between the datasets, the differential expression of genes quantified in WCT, CyT and PoP were tabulated alongside the lists of GRASPS or PoP DEGs and further sorted out by fold changes in PoP or GRASPS respectively. Two conserved blocks of 430 and 356 up- and down-regulated genes are commonly identified between the 2 translatomes, while the transcriptomes show distinct patterns with approximately half of the translatome changes not detected as differentially-expressed (**Fig. 4c**). On the other hand, sorting out the differential gene expression tabulated alongside GRASPS or PoP by fold changes in WCT highlights altered expression of two conserved blocks of 280 and 266 up- and down-regulated genes commonly detected in the translatomes but not differentially-expressed in the WCT (**Fig. 4d**). Overall, this analysis showed that the expression levels of 546 out of 786 (69%) of genes commonly detected as altered in the disease model by both translatome technologies were, however, not detected to be changed at the global transcriptome level. Reciprocally, the differential expression of genes quantified in CyT, PoP and GRASPS was tabulated alongside WCT and sorted out by fold changes in PoP or GRASPS respectively. Two blocks of 373 and 335 up- and down-regulated genes in WCT were not changed in the translatomes, indicating that 47% of differentially-expressed transcripts in the global transcriptome (708 out of 1,521 changes) are not changed in the translatomes (**Fig. 4e**). The full lists of gene IDs and fold change values in the clustered heatmaps presented in **Figs 4c-e** are provided in **Supplementary Data 5 tab2**. Taken together, this analysis validated that both translatomes were related, with a vast majority of gene expression changes not observed in the global transcriptome while reciprocally half of the RNA expression changes in the transcriptome did not occur at the translating level.

**Fig. 4.**
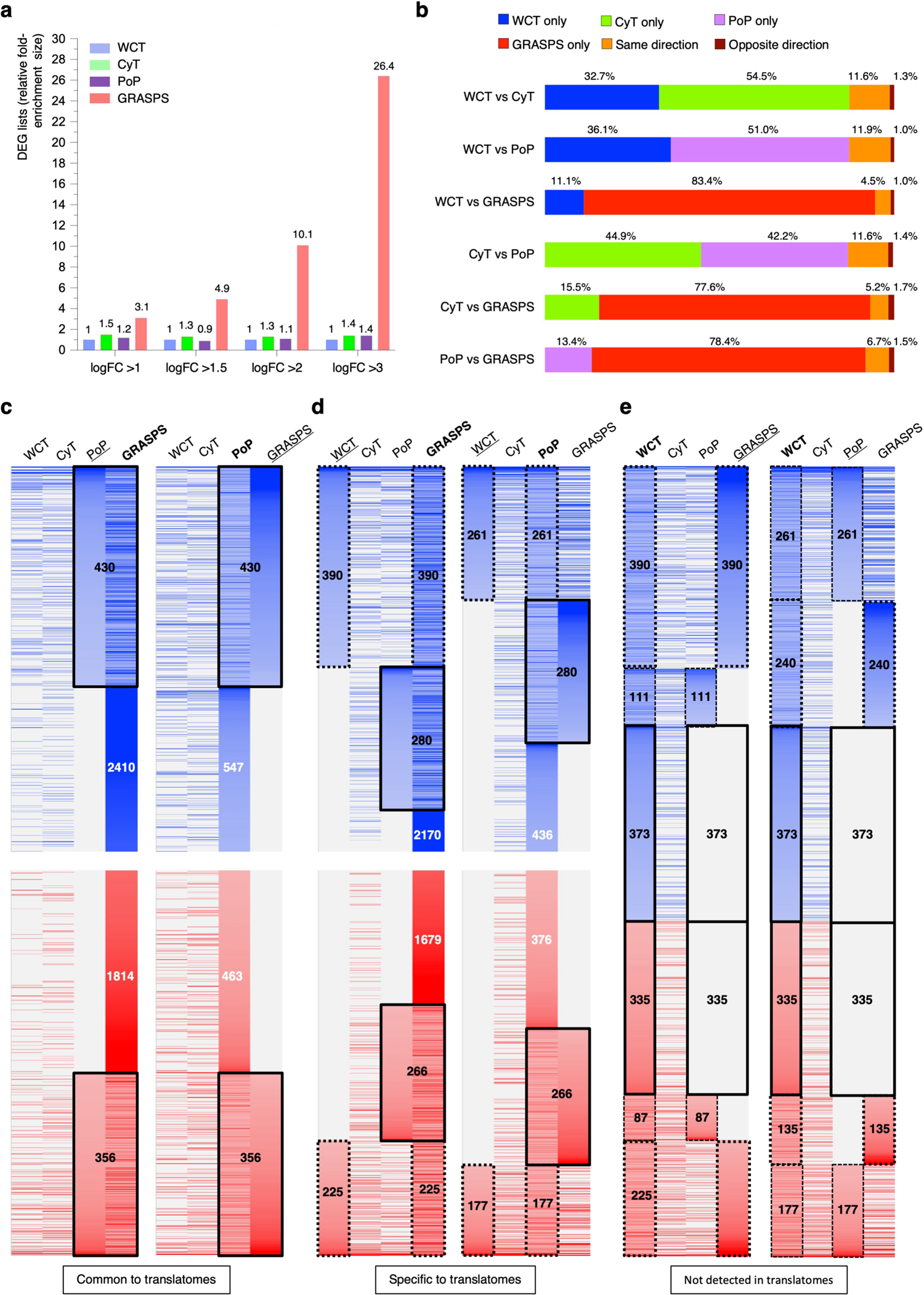
Genome-wide integration of transcriptomes and translatomes. **(a)** Bar chart representing the fold-enrichment in the number of DEGs identified in each dataset at increasing FC threshold. The size of WCT DEGs is setup as 1. **(b)** Barplots representing the percentage of uncoupling between the DEGs identified in pairwise dataset combinations. Blue, green, purple and red show DEGs specific to WCT, CyT, PoP and GRASPS respectively. Orange or brown indicate commonly-identified DEGs with same or opposite direction of changes respectively. **(c-e)** Heatmaps representing the degree of correlation between DEGs identified in the 4 datasets. They show DEGs commonly identified in the PoP and GRASPS translatomes (c), DEGs specific to the translatomes i.e. not differentially-expressed in the transcriptomes (d) and transcriptome-altered DEGs which are not changed in the translatomes (e). To generate these heatmaps, the values of fold changes quantified in the lists of DEGs were aligned/tabulated on the DEG lists identified in GRASPS and PoP (bold, c-d) or WCT (bold, e) and further sorted by fold changes on PoP and GRASPS (c, underlined), WCT (d, underlined) or GRASPS and PoP (e, underlined). A gradient of blue and red respectively indicate significantly up- and down-regulated gene expression changes for FC>2, the darker the colour the higher the FC. Grey represents genes which are not differentially-expressed. Heatmaps representing the top 750 up- and down-regulated DEGs are represented in panels c-d. Numbers indicate all DEGs counts for each list. Bold solid-line rectangles show DEGs common to translatomes (c), specific to translatomes (d) and not changed in translatomes (e). Dash-line rectangles highlight common DEGs between transcriptomes and translatomes.

### GRASPS reveals a greater number of functionally better-defined biological processes

A gene ontology (GO) investigation was performed with the protein-coding changes identified in the different datasets to further interrogate the apparent lack of correlation between translatomes and transcriptomes. Since, some transcript isoforms are up-regulated while others corresponding to the same gene are down-regulated, we performed the ontology on transcripts with either increased or decreased expression changes at gene level using the functional annotation clustering tool in DAVID (Database for Annotation, Visualization and Integrated Discovery)^58, 59^ for each of the differential expression change thresholds (**Supplementary Data 6**). Enrichment scores relating to biological processes in the most prominent clusters were plotted in heatmaps to statistically summarize the ranking of altered pathways in each of the transcriptomes, PoP and GRASPS at fold changes over 2.0, 2.8, 4.0 and 8.0 for up- and down-regulated gene expression changes (**Figs. 5a** and **5b** respectively). First of all, the same pathways are found to be enriched in either up- or down-regulated DEGs, in line with both increased or decreased expression of various gene products being involved in the regulation of a biological process, and in agreement with ALS leading to dysregulation of multiple biological processes rather than specifically affecting just one or a few from functioning. Cellular stress responses, cell death/apoptosis, DNA damage/repair, transcription, RNA splicing, cell cycle/cytoskeleton, proteolysis, autophagy and neuronal-related are known to be altered in ALS and have previously been reported in previous transcriptomics studies^60–62^. Interestingly, additional pathways relevant to ALS, but not typically identified in transcriptomics studies, are also highlighted by GRASPS. They relate to cytoplasmic stress granules, mRNA transport, cell aging or neuron death, together with others not thoroughly investigated in ALS, such as mRNA export from nucleus, mRNA stabilization/destabilization, translation initiation and tRNA biosynthesis. Moreover, this investigation confirmed that GRASPS is more sensitive, allowing detection of an increased number of biological pathways with higher confidence at higher fold changes. As expected, the functional annotation clustering combining both up- and down-regulated DEGs, except for the allowed inclusion of only 3,000 GRASPS DEGs (over 4,117), highlighted the same dysregulated pathways (**Supplementary Data 6**).

**Fig. 5.**
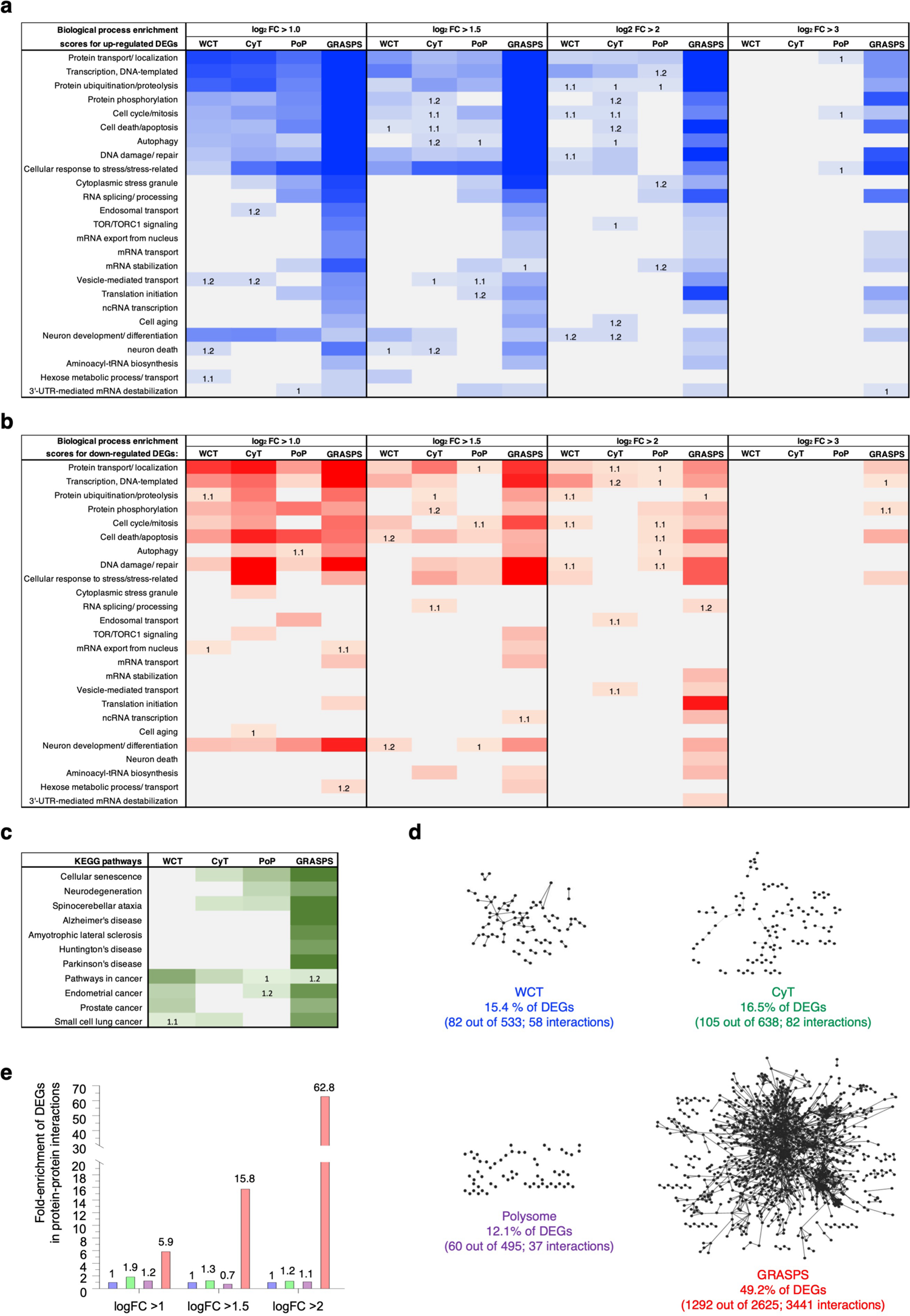
GRASPS reveals a greater number of functionally better-defined biological processes. **(a-b)** Heatmaps representing gene ontology (GO) enrichment scores (ES) for transcriptome and translatome datasets using the lists of up-regulated (a) and down-regulated (b) protein-coding DEGs filtered at various FC thresholds. The Functional Annotation Clustering tool was used in DAVID^58, 59^ based on GOTERM_ BP_FAT, GOTERM_MF_FAT, GOTERM_CC_FAT and KEGG pathways with default medium classification stringency scale. A gradient of blue and red respectively indicate significantly up- and down-regulated pathways, the darker the colour the higher the ES. Grey represents pathways not altered in the corresponding dataset. ES of 1.3 and above are considered statistically significant^58, 59^. Numbers highlight altered pathways detected with ES comprised between 1 and 1.3. **(c)** KEGG pathway analysis using the protein-coding lists of DEGs altered in the ALS-inducible cell model (FC>2). ES of 1.3 and above are statistically significant^58, 59^ and represented by a gradient of green, the darker the color the higher the ES. Numbers highlight altered pathways detected with ES comprised between 1 and 1.3. **(d)** The protein-coding lists of DEGs (log_2_FC>1.5) identified in each dataset were mapped onto the STRING protein-protein interaction database^67^ using Cytoscope with default Edge setting based on experimental protein-protein interactions. The percentage of DEGs forming functional gene regulatory networks and the numbers of interactions are indicated. See Supplementary Fig. 5 for the same investigation at various FC. **(e)** Bar chart representing fold-enrichments of DEGs mapping protein-protein interaction networks at increasing FC thresholds. Data is not presented for log_2_FC>3 as no DEG mapped to a network in the WCT, CyT and PoP datasets.

The KEGG (Kyoto Encyclopedia of Genes and Genomes)^63^ database was also interrogated to define pathways at high organism or biological level. Pathways involved in neurodegeneration and cancer are often shared in an opposite state of alteration and cancer-related terms are typically retrieved in ALS transcriptomes as in our present study (**Fig. 5c**). However, neurodegeneration is also identified by both translatomes while GRASPS strikingly showed high enrichment in neurodegenerative disease terms including amyotrophic lateral sclerosis (**Fig. 5c**), highlighting once again its higher sensitivity in the detection of relevant disease-associated pathways. The expression of genes is regulated in networks with gene products, the proteins, physically or functionally interacting with one another^64–66^. We further used STRING^67^ to functionally map our lists of DEGs for the different fold change thresholds in protein association networks based on known experimentally-evidenced protein-protein interactions. Approximately half of the DEGs identified by GRASPS for FC>2 and 2.8 strikingly matched with cellular protein networks, while a smaller proportion of DEGs with a much smaller proportion of total interactions are included in networks formed by the transcriptomes or the PoP translatome (**Fig. 5d, Supplementary** Fig. 4), showing that the higher number of expression changes quantified by GRASPS does also better correlate with gene regulatory networks and regulation/alteration of biological processes at a functional level. Interestingly, over 30% of GRASPS DEGs are still included in functional protein-association networks for FC>8 while no network is detected for WCT, CyT or PoP. Overall, GRASPS respectively shows over 5-, 15- and 60-fold enrichment in functional protein networks compared with the other datasets for fold changes >2, >2.8 and >4 respectively (**Fig. 5e**). No data is available for FC >8 since the few DEGs identified in the transcriptomes and PoP translatome are not sufficient to map onto any functional network at this high stringency of filtering (**Supplementary** Fig. 4). Taken together, our data highlight that both TDP-43 Q331K ALS-linked up-regulation and down-regulation of transcripts contribute to the same altered biological processes consistent with our current knowledge of gene expression regulation/alteration at pathway levels involving integrated modules of genetically and/or physically interacting proteins^68, 69^. Strikingly, approximately half of the differentially-expressed transcripts identified by GRASPS form gene regulatory networks.

### ALS-altered response to oxidative stress and retinoic acid regulated differentiation of neurons

To represent the identified IDs of DEGs across datasets, we generated scatter plots which compare the fold changes of transcripts commonly-altered within 2 datasets: (i) CyT versus WCT (**Fig. 6a**), (ii) PoP versus WCT or CyT (**Fig. 6b**) and (iii) GRASPS versus WCT, CyT or PoP (**Fig. 6c**). As highlighted in the volcano plots and Venn diagrams in **Figs. 3j-k**, GRASPS identified an increased number of DEGs spanning higher fold changes compared to the other datasets which identified less ALS-altered transcripts with lower fold changes. *TARDBP* mRNAs encoding TDP-43 are up-regulated in all datasets in full expectation with the tetracycline-mediated induction of TDP-43 Q331K which was validated in the ALS-inducible cell model at both RNA and protein levels (**Figs. 1a-c**). On the other hand, the Spearman correlation values confirm the broad uncoupling across datasets. The much higher number of DEGs in GRASPS also contributes to increasing this effect including when comparing to the PoP despite the two conserved blocks of 430 and 356 up- and down-regulated genes commonly identified in both translatomes (**Fig. 4c**).

**Fig. 6.**
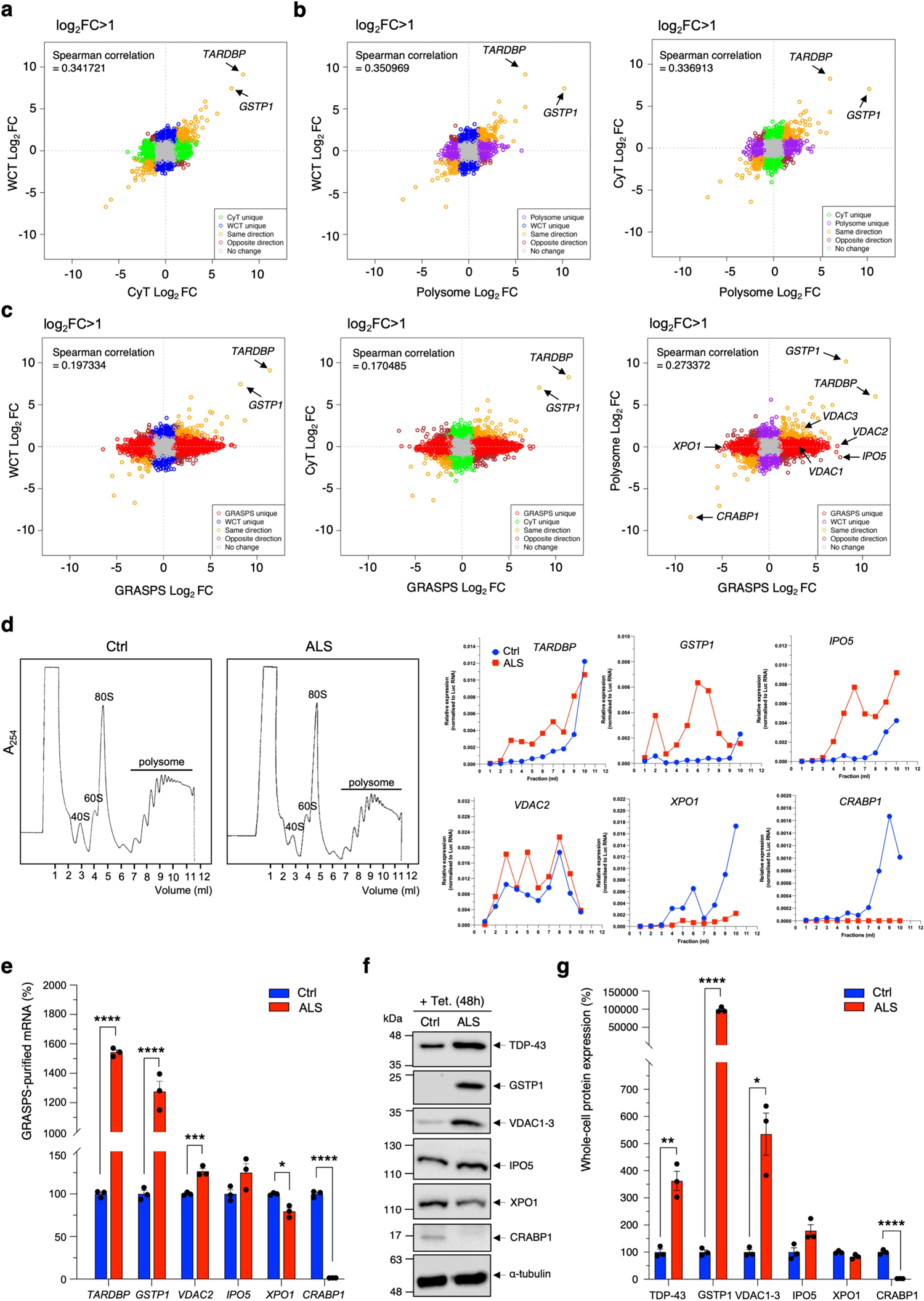
Experimental validation of key predicted DEG alterations in the human TDP-43 ALS-inducible cells. **(a-c)** Scatter plots of WCT, CyT, PoP and GRASPS log2-transformed fold changes for common transcripts in pairwise dataset comparisons. Each circle represents one transcript. Blue, green, purple and red circles show DEGs which are specific to WCT, CyT, PoP and GRASPS respectively. Orange or brown circles indicate DEGs showing same or opposite direction of changes respectively, while grey circles highlight transcripts which are not differentially-expressed in the datasets. Transcripts highlighted by arrows are either predicted to be altered in all datasets or specifically up- or down-regulated in the translatomes. They are selected for experimental validation in panels d-g. **(d)** Polysome profiling experiments in human Ctrl and TDP-43-linked ALS cell lines induced with tetracycline for 48h. Traces highlight the small 40S and large 60S ribosome subunits, the 80S initiating complexes and the actively-translating polysome fractions. Expression levels of *TARDBP* encoding TDP-43, *GSTP1*, *IPO*5, *VDAC2*, *XPO1* and *CRABP1* are quantified by qRT-PCR in each sucrose gradient fraction and normalised to a spiked luciferase RNA control for each cell lines (Ctrl in blue and ALS in blue). **(e)** GRASPS-purified mRNAs of interest (*TARDBP*, *GSTP1*, *IPO*5, *VDAC2*, *XPO1* and *CRABP1*) from tetracycline-induced (48h) Ctrl and TDP-43-linked ALS cell lines were quantified by qRT-PCR and normalised to *GAPDH* mRNA levels in three biological replicates (Mean ± SEM; unpaired multiple t tests with Holm-Šídák correction; *: p=007226, ***: p=0.001669; ****: p<0.0001, N=3). **(f)** Protein extracts from tetracycline-induced (48h) Ctrl and TDP-43-linked ALS cell lines were analysed by western immunoblotting probed with antibodies against TDP-43, GSTP1, VDAC1-3, IPO5, XPO1, CRABP1 and loading control α-tubulin. **(g)** Quantification of protein expression levels in panel (f) following normalization to α-tubulin in three biological replicates (Mean ± SEM; unpaired multiple t tests with Holm-Šídák correction; *: p=0.005108, **: p=0.002098, ****: p<0.0001, N=3).

*GSTP1* (Glutathione S-Transferase Pi 1), one of the most up-regulated transcripts at total, cytoplasmic and translating levels, encodes a protein counteracting oxidative stress but also associated with a role in susceptibility to cancers and neurodegenerative diseases^70, 71^. On the other hand, the expression levels of the most down-regulated translating transcript, *CRABP1* (Cellular Retinoic Acid Binding Protein 1), recently reported to be decreased in ALS and motor neuron degeneration^72, 73^, are not detected as differentially expressed in the whole-cell or cytoplasmic transcriptomes. The genome-wide investigation also predicted that the translation but not the total or cytoplasmic expression levels of *VDAC1-3* (voltage dependent anion channel) and *IPO5* (importin 5) transcripts are increased, while the translation of *XPO1* (exportin 1, also known as CRM1 for chromosomal region maintenance 1) is reduced in the ALS-inducible cell model. Interestingly, the nucleocytoplasmic transport pathway which involves XPO1 and IPO5 is affected in neuronal aging^74^ while overexpression of VDAC1 was recently reported in an Alzheimer’s disease model of neurodegeneration^75^; while VDACs dysfunction has been linked with carcinogenesis and neurodegenerative diseases including ALS^76^.

To experimentally validate these bioinformatics-predicted findings, particularly those showing altered translation of specific transcripts without changes in their whole-cell or cytoplasmic expression, we performed polysome profiling experiments with qRT-PCR quantification of transcripts of interest as an established gold-standard methodology to investigate actively translated RNA molecules in the 80S translation initiation complexes and the translating polysomal fractions of tetracycline-induced control and ALS cell lines (**Fig. 6d**). Accordingly, we observed that the translation of *TARDBP*, *GSTP1* and *IPO5* transcripts is higher in the ALS-induced compared to the control cells while the translation of *VDAC2*, *XPO1* and *CRABP1* mRNAs is indeed reduced in the ALS cell model. We further used qRT-PCR to directly quantify these transcripts in GRASPS-purified ribosomes isolated from the tetracycline-induced control and ALS cell lines and confirmed the previous polysome profiling validation and bioinformatics prediction (**Fig. 6e**). Consequently, we also found that the expression levels of TDP-43, GSTP1 and IPO5 proteins are increased while those of VDAC, XPO1 and CRABP1 are reduced in the ALS-inducible cell model (**Fig. 6f**, quantification in **Fig. 6g**), thus providing functional relevance to the predicted alteration of translatomes and gene expression changes identified by GRASPS.

## Discussion

To our knowledge, this study compares for the first-time head-to-head genome-wide abundance of whole-cell (total), cytoplasmic and translating RNAs using a stable isogenic inducible human cell model which allows for low and time-controllable induction of the TDP43 Q331K-linked ALS mutant protein to identify the most upstream disease-altered expression of genes in a 48h window of expression. It also provides high-depth RNA-seq datasets which can be used by the scientific community as a resource for multiple other comparative biological and computational investigations. While it is known that transcriptomes and proteomes significantly differ, we also found that the cytoplasmic expression levels of RNAs also do not correlate with the abundance of translating transcripts associated with ribosomes. Overall, this suggests that the proportion of RNA translated in a cell represents only a small proportion of the total or cytoplasmic amounts of RNA molecules. Strikingly, approximately half of the gene expression alterations identified by transcriptomics are not dysregulated in the translatomes while, reciprocally, the levels of 69% of transcripts found to be commonly affected in both translatome methodologies are not changed in the whole-cell transcriptome.

Despite broad uncoupling in the differentially-expressed transcript IDs characterized among the various datasets, it is worth noting that the altered pathways are common with the identification of biological processes known to be altered in ALS tissues or patient-derived neurons. These include cell death/apoptosis, responses to cellular stress, DNA damage/repair and cell cycle, altered RNA metabolism, dysregulated protein homeostasis and neuronal-related damages among others^60–62^. In addition to validation of this stable TDP-43 ALS-inducible HEK cell line as a relevant model of disease, our findings show that the widespread dysregulation of gene expression occurs in a fast process, within 48 hours, and simultaneously affects multiple cellular processes with altered expression of approximately 1,400 and 4,400 genes in the global transcriptome and the GRASPS translatome respectively. Other studies including the ribosome profiling investigation of TDP-43 A315T ALS-linked translatome reported the differential expression of 26 and 51 genes in motor neuron-like cells and primary cultures of cortical neurons respectively^77^ while TRAP translatomes identified 616 specific gene expression changes in a TDP-43 G298S ALS *Drosophila* model together with another 1,892 DEGs due to TDP-43 overexpression^78^. On the other hand, TRAP investigations have identified translatomes in the spinal cord motor neurons of a TDP-43 A315T ALS mouse model with specific alteration of 1,061 DEGs in 14-week old symptomatic mice^79^ while 28 gene expression changes were reported in 10-month old mice^80^.

Here, we also present a novel translatome methodology, GRASPS, to bypass: (i) the very limited throughput and requirement for specific equipment and expert skills that restrain the universal investigation of translatomes; (ii) the addition of artefact-causing chemical stressors; (iii) the use of sucrose gradients which requires delicate manipulation and hinders accurate fractionation and reproducibility; (iv) or the necessary tagging of ribosomal subunits and time-consuming generation of transgenic cell lines or animal models^9, 19^. Strikingly, GRASPS led to the detection of approximately 3.5 times more gene expression changes with higher fold changes and lower p-values compared to the WCT/ CyT transcriptomes and the polysome profiling method. The simplicity and faster purification of ribosome-associated RNAs is attributed to the higher reproducibility and improved statistical power of GRASPS, in contrast to more complex protocols involving sucrose gradient fractionation or affinity purification. Moreover, our data shows that GRASPS identified large blocks of up- and down-regulated gene expression changes (430 and 356 respectively) which are commonly conserved with actively translated RNAs characterized in the polysome fractions, thus providing a genome-wide level validation for the new methodology. In addition, we show that GRASPS-altered gene expression changes map to better-defined dysregulated biological processes at a functional protein-association level with approximately half of DEGs forming known gene regulatory networks. Interestingly, additional pathways relevant to ALS are also highlighted by GRASPS such as cytoplasmic stress granules, mRNA transport, mRNA export from nucleus, cell aging and neuronal death. Exceeding our best expectations in the KEGG pathway investigation, GRASPS was the only method to retrieve the term “amyotrophic lateral sclerosis” in a human embryonic kidney cell model of ALS. We and others applied GRASPS to other mammalian cell types including mouse motor-neuron like NSC34 cells and patient-derived astrocytes or neurons. Future work will aim to further test and optimize this technology in animal or human post-mortem tissues.

## Methods

### Cell culture

TDP-43 Q331K (ALS) and sham control (Ctrl) HEK293T Flp-In cell lines were generated according to the manufacturer’s instructions from Invitrogen. In brief, HEK FRT host cells harboring the Flp-In™ T-REx system were maintained in culture medium (DMEM-high glucose (Sigma) medium, 10% tetracycline-free FBS (Biosera) and 1% penicillin/ streptomycin (Lonza)) with selection antibiotics blasticidin S (15 μg/ml, Calbiochem) and zeocin (100 μg/ml, Invitrogen). To establish the tetracycline-inducible TDP-43 Q331K ALS cell line or control cell lines, HEK FRT cells were plated in 10-cm dish with culture medium containing only blasticidin S following co-transfection with Flp recombinase expressed plasmid pPGKFLPobpA and tetracycline-responsive plasmid pcDNA5FRT/TO containing TDP-43 Q331K or pcDNA5FRT/TO only as isogenic sham control in the ratio 6:4 using PEI (polyethyleneimine; Sigma) transfection method. The fresh medium containing blasticidin was changed 24 hours post-transfection. After another 24 hours, the transfected cells were split and plated into three 10-cm dishes in culture medium containing selection antibiotics blasticidin S (15 μg/ml, Calbiochem) and hygromycin B (100 μg/ml, Invitrogen). The selection medium was changed every 3 days until foci were identified. Hygromycin resistant foci were expanded before verifying that the pcDNA5/FRT/TO construct had integrated into the FRT site by testing each clone for zeocin sensitivity. The picked colonies for control sham and TDP-43 Q331K ALS were later maintained in T75 flask (2×10^6^ cells) in culture medium containing blasticidin S (15 μg/ml) and hygromycin B (100 μg/ml). Cells were plated on 10 cm dishes (2×10^6^ cells), 6 well plates (2×10^5^ cells/well) or 24 well plates (5×10^4^ cells/well) with or without coverslip and induced with tetracycline (10 μg/ml, Invitrogen) for 48 hours. Tetracycline-containing medium was changed every 48 hours if a longer induction time was needed.

### Cell growth curve

HEK293T Flp-In cells (1 x 10^6^ cells) were plated in 10 cm dishes in the presence or absence of tetracycline. At three-day intervals the cells were trypsinised and resuspended in 5 ml of media. The cells were then counted using a haemocytometer and the total number of cells in each plate calculated. After counting, 1 x 10^6^ cells were returned to the 10 cm plate and given fresh media containing tetracycline where required. This was repeated for the duration of the growth curve.

### Total and cytoplasmic RNA fractionation and extraction

HEK Flp-In cell models were grown in 10-cm dishes for 24 hours before induction with tetracycline. Cells were lysed post tetracycline induction when an approximate confluency of 70-80 % had been achieved. Total RNAs were collected from a 10-cm dish directly in 400 μl of 1x Reporter lysis buffer (Promega) containing 0.16 U/μl Ribosafe RNase inhibitors (Bioline), 2 mM PMSF and cOmplete™ EDTA-free Protease Inhibitor Cocktail and lysed for 10 min on ice before refrigerated centrifugation at 17,000g for 5 min. The supernatant was collected.

For cytoplasmic fractions, cells were lifted from a 10-cm dish in PBS and centrifuged at 400g for 5 min, 4°C to pellet cells prior to fractionation. Cell pellets were quickly washed with hypotonic lysis buffer (10 mM HEPES pH 7.9, 1.5 mM MgCl2, 10 mM KCl, 0.5 mM DTT) prior to lysing cells in 400 μl hypotonic lysis buffer containing 0.16 U/μl RNasin, 2 mM PMSF and cOmplete™ EDTA-free Protease Inhibitor Cocktail. Cells were resuspended gently using a cut P1000 tip and left on ice for 10 mins. The cytoplasmic fraction was collected after differential centrifugation at 4°C (3 min at 1500g, 8 min at 3500g, and 1 min at 17,000g). PureZole (Bio-Rad) was added to total fractions and cytoplasmic fractions and RNA was extracted using Direct Zol RNA Miniprep Plus (Zymo Research) for RNA sequencing or using the standard chloroform and ethanol protocol for cDNA synthesis and RT-qPCR.

### Polysome profiling

HEK Flp-In cells were plated at a cell density of 2.5 x 10^6^ in T175, with four T175 flasks per cell line. After 48 h of tetracycline induction, cells were treated with cycloheximide (100 µg/ml) on ice for 30 min to stall the ribosome on the mRNAs. Cells were scraped in the cell medium and collected into a 50 ml falcon tube. After refrigerated centrifugation at 500 x g for 5 min, the medium was discarded. Cell pellets were resuspended with 7 ml polysome buffer (20 mM HEPES pH7.4, 100 mM KOAc, 2 mM MgOAc, 0.5 mM DTT, 100 μg/ml cycloheximide) and transferred to a 15 ml falcon tube. Cells were pelleted by refrigerated centrifugation at 500 g for 5 min. The supernatant was discarded and cell pellets were resuspended in 800 µl polysome lysis buffer before transfer to 1.5 ml microtubes. Cells were pelleted by refrigerated centrifugation at 500 g for 5 min. After complete removal of the supernatant, the cells were lysed with polysome lysis buffer containing 0.5% Triton X-100, 0.16 U/μl RNase inhibitor (Bioline), 2 mM PMSF and cOmplete™ EDTA-free Protease Inhibitor Cocktail. Cell lysates were passed through a 25G needle 10 times and left on ice for 10 min. Cell extracts were transferred to fresh 1.5 ml microtubes and centrifuged at 10,000 rpm, 4°C for 15 min. The OD_260_ unit of cell lysate was determined using a Nano drop. Cell extracts (∼12 OD_260_ unit) were loaded onto 15-50% sucrose gradient and ribosomes pelleted by ultracentrifugation (Th-641 swing-out rotor, Sorvall WX) at 80,000 rpm, 4°C for 2 h. Ribosome fractions (0.5 ml/ fraction) were collected into PureZole containing spiking luciferase RNA using a ISCO gradient collection and fractionation system which continuously measures the A_254_ to provide the polysome traces. RNAs were extracted from each fraction using Direct Zol RNA Miniprep Plus (Zymo Research) for RNA sequencing.

### Genome-wide RNA Analysis of Stalled Protein Synthesis (GRASPS)

HEK Flp-In sham and TDP-43 Q331K cells were induced with 10 μg/ml tetracycline for 48 h after plating cells in four 10 cm plates per line for 48 h (2 x 10^6^ cells per plate). Cells were washed with cold DEPC-treated PBS on ice and collected using a cell scraper in 2 ml cold DEPC-treated PBS. Cells were pelleted by centrifugation at 400 x g for 5 min at 4°C. The PBS was discarded, and the leftover of PBS was removed by carefully adding 200 μl GRASPS buffer A (250 mM sucrose, 5 mM KCl, 50 mM Tris-HCl pH 7.4) without disturbing the pellet. A volume of GRASPS buffer A1 containing 0.16 U/μl RNase inhibitor (Bioline), 2 mM PMSF and cOmplete™ EDTA-free Protease Inhibitor Cocktail corresponding to approximately three times the volume of the cell pellet (600 μl) was added to resuspend the pellet by pipetting up and down with a cut p1000 tip. Cells were lysed by pipetting up and down 56 μl 10% NP-40 with a cut p200 tip to reach a final concentration of 0.7% v/v. The lysis is achieved on ice for 10 min swirling gently the tubes from time to time. The lysates were transferred to one well of a 6-well plate for each condition and UV-irradiated at 0.3 J/cm^2^ on ice. Nuclei are pelleted by centrifugation at 750 x g, 4°C for 10 min and the supernatants were carefully transferred to prechilled RNase-free tubes before centrifugation at 12,500 x g, 4°C for 10 min to pellet the mitochondria. Typically, 480-500 μl of supernatants, containing the organelle-free cytoplasmic fraction, were transferred to fresh prechilled tubes. All conditions were adjusted with GRASPS buffer A1 to 500 μl and 71 μl 4 M KCl solution was added to a final concentration of 0.5 M KCl. 429 μl GRASPS buffer B (250 mM sucrose, 500 mM KCl, 50 mM Tris-HCl pH 7.4, 5 mM MgCl_2_) is added to top up to 1 ml before carefully loading onto 1 ml sucrose cushions (1 M sucrose, 5 mM MgCl_2_, 50 mM Tris-HCl pH 7.4) in cold TLA100 centrifuge tubes. Ribosomes were pelleted by ultracentrifugation at 250,000 x g, 4°C for 2 h. After discarding the supernatants, the leftover of sucrose was washed carefully by adding 150 μl cold DEPC-treated water and removed immediately.

The glossy pellets of ribosomes were resuspended in 250 μl cold ribosome resuspension buffer (50 mM HEPES pH 7.9, 150 mM NaCl, 1 mM DTT, 1 mM EDTA) and transferred to fresh RNAse-free tubes. Proteinase K was added to a final concentration of 100 μg/ml (1.25 μl of stock at 20 mg/ml) with 0.16 U/μl Ribosafe RNase inhibitor (Bioline). The reactions were incubated at 37°C for 30 min after pulse vortex to digest the ribosomal proteins. The proteinase K was inactivated by adding 5 μl of 0.5 M EDTA (final concentration 10 mM) and 4.2 μl of 3 M NaAc (final concentration 50 mM). PureZole (Bio-Rad) was added to each GRASPS-purified RNA fraction and RNA was extracted using Direct Zol RNA Miniprep Plus columns (Zymo Research) for RNA sequencing or a standard chloroform and ethanol precipitation for cDNA synthesis and RT-qPCR.

### Liquid chromatography tandem mass spectrometry (LC-MS/MS)

Trypsin-digested peptides were analyzed by nano-HPLC (UltiMate 3000 HPLC system; Thermo, Hemel Hempstead, UK) coupled to an amaZon ETD MS ion trap spectrometer (Bruker Daltonics, Bremen, Germany) using a nano-ESI spray. The nano-HPLC system and the ion trap spectrometer were controlled using Bruker Compass HyStar v3.2-SR2 software. The liquid chromatography system comprised of a reversed-phase precolumn (LC Packings, Dionex) for sample desalting and a PepMap 100 reversed-phase C18 column (75 μm by 15 cm; Thermo) for peptide fractionation. The flow rate for precolumn loading was 30 μl/min of loading buffer (97% [vol/vol] ACN and 0.1% [vol/vol] TFA). Peptides were analyzed at a flow rate of 300 nl/min and separated by gradient elution using buffer A (3% [vol/vol] ACN and 0.1% [vol/vol] FA) and buffer B (97% ACN and 0.1% FA [vol/vol]) as follows: 4% buffer B (0 to 5 min), 5 to 38% buffer B (5 to 65 min), 38 to 90% buffer B (65 to 68 min), and 90% buffer B (68 to 73 min), followed by re-equilibration at 4% buffer B. The electrospray was operated in positive-ion mode with a 4,500-V spray voltage, 10-lb/in2 gas pressure, and 150°C dry gas. The end plate offset of the mass spectrometer was set to −500 V and data acquisition using standard method Proteomics Auto MSMS.

#### Protein database searching

Mass spectra were first converted to Mascot Generic Files using scripts provided by Bruker (Bremen, Germany). The Mascot server v2.5 database search engine^81^ was used for peptide and protein identification. Databases used were the cRAP (https://www.thegpm.org/crap/) 116 entries (downloaded on 26^th^ February 2024) and UniProt Homo sapiens proteome database (82,485 entries downloaded on 26^th^ February 2024). The search parameters were trypsin/P digest, 2 missed cleavages, 1.2 Da precursor tolerance, 0.6 Da fragment tolerance, carbamidomethyl cysteine modification set to fixed, and methionine oxidation and N-Acetyl modifications as variable. Charge states included +2, + 3, and +4. Data was searched using Mascot’s integrated reversed decoy database method to calculate FDR. A Mascot-integrated decoy database search calculated an FDR of 1%. The Percolator algorithm in MASCOT was applied to improve the distinction between correct and incorrect spectral matching^82^. The resulting protein list is available in Supplementary Data 4.

### Purification of poly(A)+ mRNA extracted from GRASPS samples

NEB Next® Poly(A)+ mRNA Magnetic Isolation Module (New England BioLabs Inc.) was used to purify mRNA based upon on the coupling of oligo d(T)25 to 1 μm paramagnetic beads which were then used as the solid support for the direct binding of poly(A)+ RNA. Beads were separated from the supernatant using a magnetic rack. The integrity of total RNA was assessed using an RNA Pico Chip (Agilent Technlogies, Inc.). <u>The purification</u> <u>protocol for the poly(A)+ RNA was optimized from the manufacturer (NEB) as ribosomes have very</u> <u>high content of rRNAs</u>. 20 μl of Oligo d(T)25 beads were placed in 0.2 ml PCR tube and washed twice with 100 μl of 2 X RNA binding buffer to remove the supernatant. A 100 μl volume of 2 X RNA binding buffer was then added to the beads. Total RNA was diluted with DEPC-treated water to a final volume of 50 μl and added to the magnetic beads in 50 μl RNA binding buffer. Samples were incubated at 65°C for 5 min and placed on ice for 2 min to denature RNA and facilitate the binding of poly(A)+ RNA to the beads. Samples were incubated at RT for 5 min to allow RNA to bind to the beads. The tubes were then placed on the magnetic rack for 2 min to separate poly(A)+ RNA bound to the beads from the solution. The supernatant was taken and kept. At this point the protocol was optimized, and an additional binding step carried out which increased the yield of ribosome-associated poly(A)+ RNA.

The beads were washed twice with 200 μl wash buffer, pipetting the volume up and down 6 times to ensure thorough mixing and placing on the rack for 2 min to ensure proper separation of unbound RNA. The beads were then stored on ice. The saved supernatant (from the initial binding) was then incubated at 65°C for 5 min and placing on ice for 2 min. The binding of poly(A) + RNA was then repeated as above with the beads stored on ice. The beads were then washed twice with 200 μl wash buffer, pipetting the volume up and down 6 times to ensure thorough mixing and placed on the rack for 2 min to ensure proper separation of unbound RNA. After ensuring total removal of wash buffer, 50 μl elution buffer was added to the beads and mixed well by pipetting. Tubes were placed at 80°C for 2 min then placed at RT immediately to elute the poly(A) + RNA from the beads. 50 μl of 2 X RNA binding buffer was added to each sample pipetting the volume up and down 6 times to ensure thorough mixing to allow RNA to bind to the beads. Samples were incubated at RT for 5 min with agitation every few minutes during the incubation. The tubes were allowed to stand on the magnetic rack for 2 min before removing the supernatant and washing the beads twice with 200 μl wash buffer, pipetting the volume up and down 6 times to ensure thorough mixing and placed on the rack for 2 min to ensure proper separation of unbound RNA. The mRNA was eluted by adding 20 μl of elution buffer to the beads and incubating at 80°C for 2 min before immediately putting the samples onto the magnetic rack for 2 min. Purified mRNA was transferred to a clean nuclease-free PCR tube before the yield and size distribution was assessed using an RNA Pico Chip (Agilent Technologies, Inc.).

### Next generation RNA sequencing and quantification of transcript abundance

Total RNAs and polyA(+)-enriched RNA samples were sent to the Centre for Genomic Research at the University of Liverpool for the preparation of dual-indexed strand-specific RNA-seq libraries (projects LIMS4821 and LIMS20433). Paired-end multiplexed sequencing runs were performed on Illumina Hi-Seq platform (2x 100bp, generating data of in excess of 120M clusters per lane; WCT, CyT, GRASPS) or Illumina HiSeq 4000 (2×150 bp sequencing, generating data from >280 M clusters per lane; PoP). Sequencing reads were aligned on GRCh38 (hg38) using Bowtie^50^. RNA expression levels were quantified using BitSeq^51^ and Limma^52^. For the statistical analysis, differentially-expressed (DE) transcripts were filtered for log2FC>1, p-value p<0.05 and false discovery rate FDR<0.2.

### Quantitative RT-PCR (qRT-PCR)

Following RNA extraction and quantification, 2 μg of DNase I (Roche) treated RNA was converted to cDNA using BioScript Reverse Transcriptase (Bioline) with (dN)_6_ random primers or dT_18_ primers as described in^40, 42, 43^. qRT-PCR reactions were performed in duplicate using the Brilliant III Ultra-Fast SYBR Green QPCR Master Mix (Agilent Technologies) on a C1000 Touch™ thermos Cycler using the CFX96™ Real-Time System (BioRAD) using an initial denaturation step, 45 cycles of amplification (95°C for 30 s; 60°C for 30 s; 72°C for 1 min) prior to recording melting curves. qRT–PCR data was analyzed using CFX Manager™ software (Version 3.1) (BioRAD) and GraphPad Prism (Version 9.0). Primers used in this study are provided in Supplementary Table 1.

### SDS-PAGE electrophoresis and Western Blot

HEK293T Flp-In cell lines, Ctrl and ALS, were plated in 6-well plate (5×10^4^/well) and induced with or without tetracycline (10 μg/ml) for 48 hours prior to harvest. Cells were washed in ice-cold PBS and lysed in lysis buffer (50 mM HEPES pH 7.5, 150 mM NaCl, 1 mM DTT, 0.5 % Triton X-100, 1 mM EDTA) supplemented with 2 mM phenylmethylsulfonyl fluoride (PMSF, Sigma) and cOmplete™, EDTA-free Protease Inhibitor Cocktail (Roche) and left on ice for 10 mins. Proteins were collected by centrifugation at 13,300 rpm for 5 mins in the refrigerated centrifuge. Protein concentration was quantified using Bradford reagent (Bio-Rad). Proteins were separated using SDS-PAGE, electroblotted onto nitrocellulose membrane and probed with indicated primary antibodies. Target protein signals were amplified with the relevant secondary antibody conjugated with HRP (Horseradish peroxidase) and chemiluminescence signals were captured by LI-COR Odyssey imaging system and intensity was quantified by Image Studio Lite software. Antibodies used in this study are described in Supplementary Table 2.

### Immuno-fluorescence microscopy

HEK293T Flp-In cells (5×10^4^/well) were plated on coverslip in 24-well plate and induced with tetracycline for 48 h. Cells were fixed and permeabilized with 4 % paraformaldehyde (PFA) containing 0.2 % Triton-X at room temperature for 20 mins. After blocking in 2 % w/v bovine serum albumin (BSA, Sigma) in PBS at room temperature for 1 hour, cells were incubated in primary antibody at room temperature for 1 hour. Cells were washed three times with PBS prior to incubation with the relevant fluorophore-labelled conjugated secondary antibody at room temperature for 30 mins. The cells were washed with PBS three times and incubated with Hoechst (Sigma 33258) for 10 min at room temperature. The coverslips were then mounted onto glass slides using fluorescent mounting medium (Dako). Images were captured using fluorescence microscopy (Nikon).

### Statistics and reproducibility

Two-way ANOVA (analysis of variance) with the recommended Tukey’s correction for multiple comparison tests (GraphPad Prism) were used for all normally-distributed data which involved more than 2 experimental conditions. Quantification of qRT-PCR and western immunoblot comparing various primers or proteins in 2 cell lines used unpaired multiple t tests with the recommended Holm-Šídák method (assuming both groups have same standard deviation of the mean). At least 3 independent biological replicate experiments were generated for statistical analysis. Data were plotted using GraphPad Prism version 9. Significance is indicated as follows; NS: non-significant, p`≥`0.05; *: p<0.05; **: p<0.01; ***: p<0.001; ****: p<0.0001.

### Ethics statement

Ethical approval was not required for this study.

## Acknowledgments

G.M.H. acknowledges support from a Faculty of Medicine, Health and Dentistry (FMDH) Research and Innovation Award (312366) from the University of Sheffield, the Royal Society research grant RG140690, the Medical Research Council (MRC) New Investigator research grant MR/R024162/1 and the Biotechnology and Biological Sciences Research Council (BBSRC) grant BB/S005277/1 over the course of this study. Y.H.L. was initially supported by a Postdoctoral Research Abroad Program sponsored by the Taiwanese Ministry of Science and Technology (105-2917-I-564-070). JED was supported by the University of Sheffield Moody Family Endowment (CPW) and the Motor Neurone Disease Association (grant Hautbergue/Mar16/900-790). S.R.M. was supported by a Faculty PhD studentship funded by the University of Sheffield. The research of M.R.G. was conducted within the framework of the Basic Research Program at the National Research University Higher School of Economics (HSE). P.J.S. is supported as a National Institute for Health Research Senior Investigator and by the NIHR Sheffield Biomedical Research Centre. Data generation and processing of raw RNA sequencing reads were carried out by the Centre for Genomic Research which is based at the University of Liverpool (projects LIMS4821 and LIMS20433).

## Authors contributions

G.M.H. funded, designed and supervised the overall study with support from M.M. and P.J.S. J.E.D and Y.H.L. respectively developed and validated the GRASPS technology. Y.H.L. further performed all the experimental validation including polysome profiling experiments with support from S.R.M. and L.M.C in G.M.H.’s group and K.L. in S.G.C.’s lab. J.E.D and Y.H.L. characterized the HEK TDP-43 Q331K Flp-IN cell line established by A.H. in P.J.S.’s group. C.A.E. and M.J.D. characterized purified ribosomes by mass spectrometry. M.J.W. contributed to the early phase of GRASPS development. L.C. and M.M. performed the genomic alignment of RNA-seq reads and the quantification of transcript abundance and differential expression. M.M. and Y.H.L. produced the R codes for the volcano and scatter plots. M.M., L.C., Y.H.L., M.R.G, I.L., J.C.K., J.R.H. and G.M.H. contributed to the bioinformatics analysis and the generation of tables and diagrams. G.M.H. and Y.H.L. wrote the first draft of the article and prepared the figures. All authors contributed to editing the manuscript and approved the submitted version.

## Additional information

**Supplementary information** is available.

**Correspondence and requests for materials** should be addressed to Guillaume M. Hautbergue. Reagents will be shared under a Material Transfer Agreement in accordance with the University of Sheffield Tech Transfer office policy.

## Supplementary Information

**Supplementary Fig. 1.**
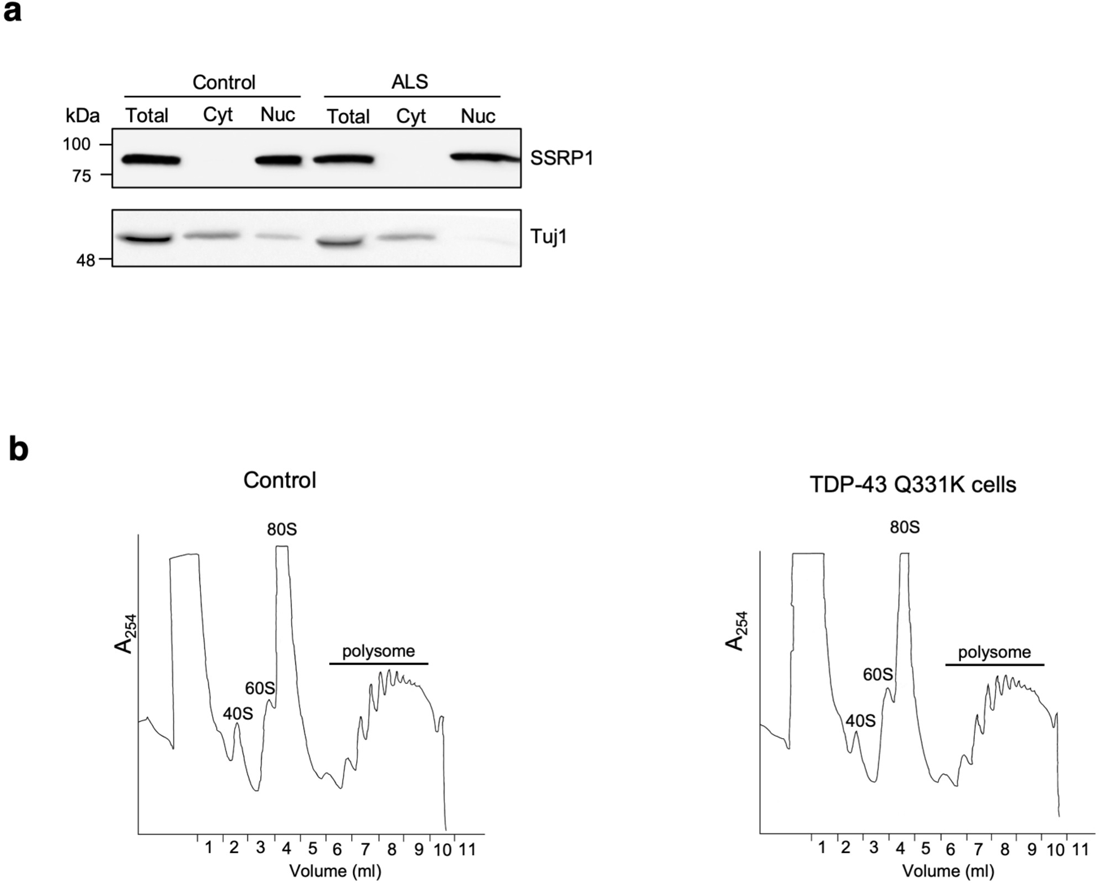
Generating a TDP-43-linked ALS-inducible stable cell model for the genome-wide identification of altered expressed and translating RNAs. (a) Protein extracts from tetracycline-induced (48h) Ctrl and TDP-43-linked ALS cell lines were analyzed by western immunoblotting probed with antibodies against nuclear chromatin-associated SSRP1 protein and cytoskeleton marker Tuj1. (b) UV traces corresponding to the polysome sucrose gradient fractionation in tetracycline-induced (48h) Ctrl and TDP-43-linked ALS cell lines. RNA was extracted from fractions 6-9, which contain actively translating polysomes (4 ml), prior to precipitation and resuspension for preparation of RNA-seq libraries.

**Supplementary Fig. 2.**
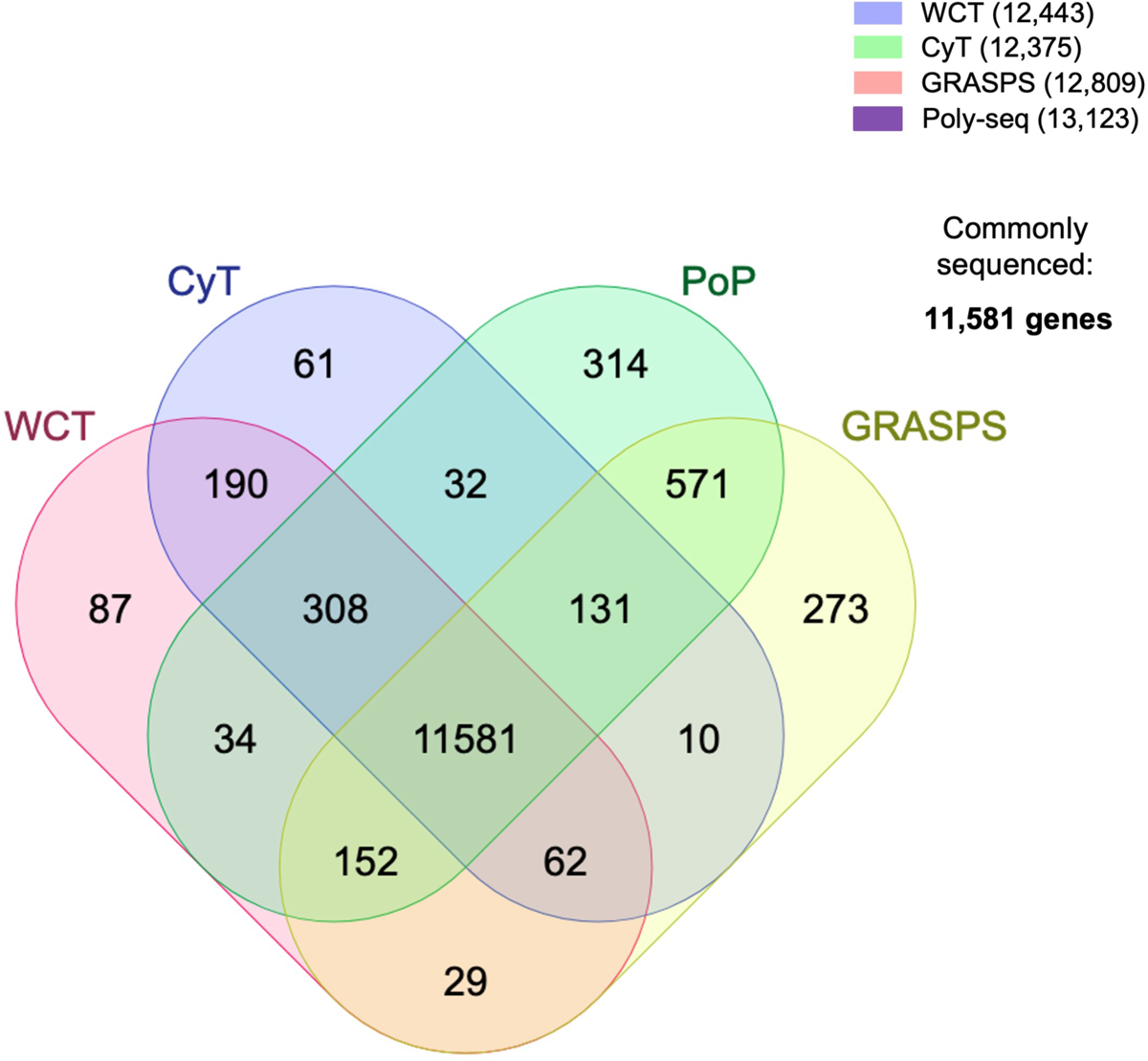
Venn diagram comparing sequenced genes across WCT, CyT, PoP and GRASPS. The Venn diagram was made using the multiple comparators online tool (https://molbiotools.com/listcompare.php).

**Supplementary Fig. 3.**
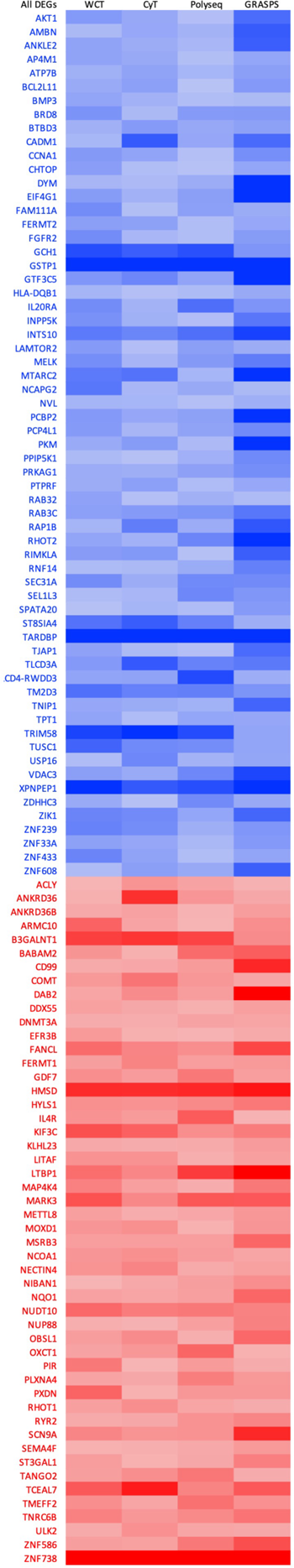
Heatmap highlighting common differentially-expressed protein-coding genes across all datasets. Genes down-regulated and up-regulated in the TDP-43 ALS-inducible cells are labelled using a gradient of red and blue colors respectively (darker the color higher the altered fold change of expression). Detailed list of genes and fold change values are provided in Supplementary Data 5 tab 1).

**Supplementary Fig. 4.**
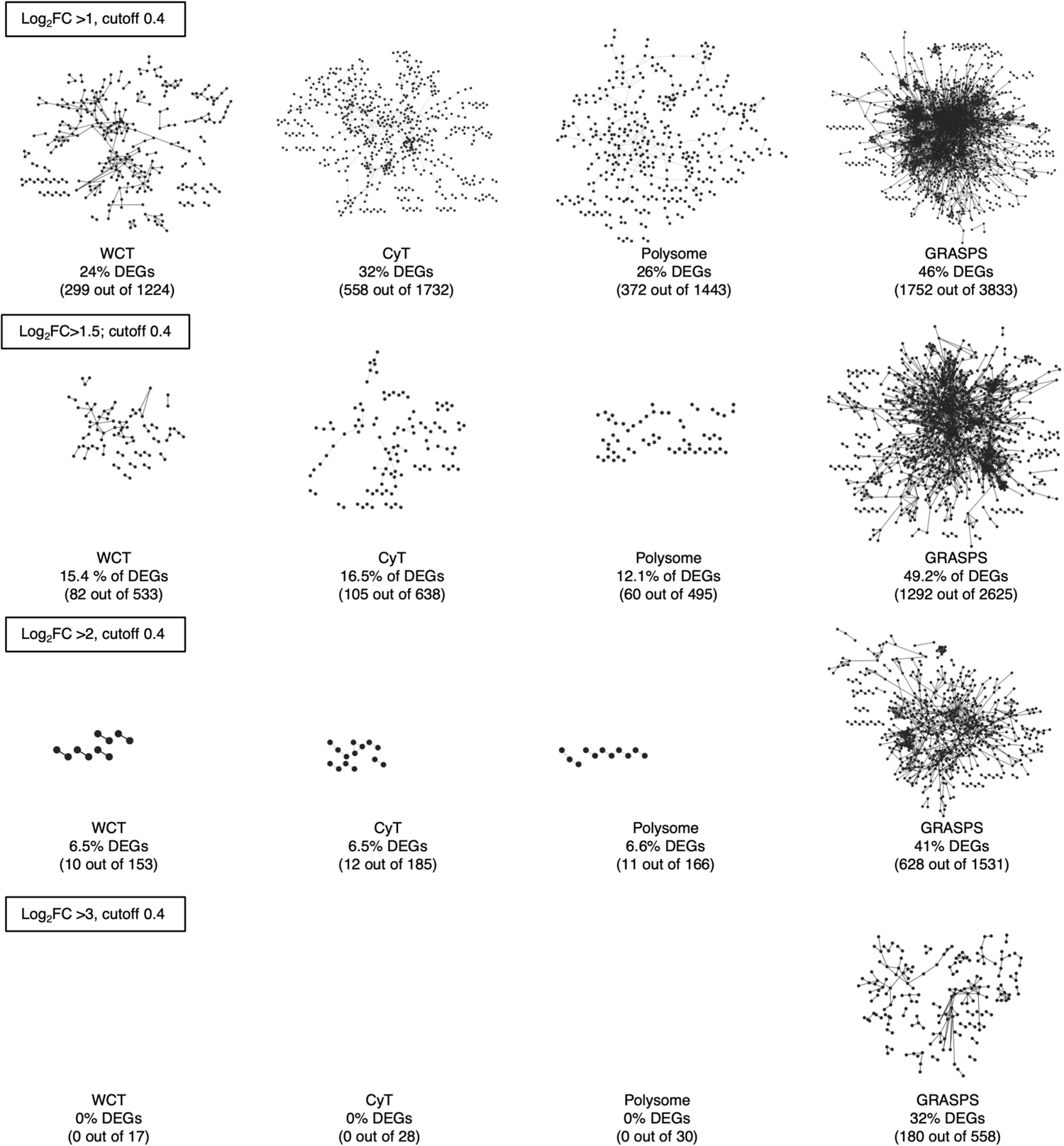
Functional gene regulatory network mapping of DEGs identified in WCT, CyT, PoP and GRASPS. Lists of protein-coding DEGs from each dataset were analysed at various FC thresholds using the String physical network database (version11.5). Identified DEGs were mapped onto functional human protein-protein interaction networks using known protein-protein interactions based on “experimental” data and default confidence Edge settings (cutoff 0.4) in the Cytoscape software environment. DEGs mapped to known gene regulatory networks are represented by solid circles while the interactions between DEGs are depicted with solid lines. The percentage of DEGs forming functional gene regulatory networks and the numbers of interactions are indicated.

---

**Supplementary Table 1.**
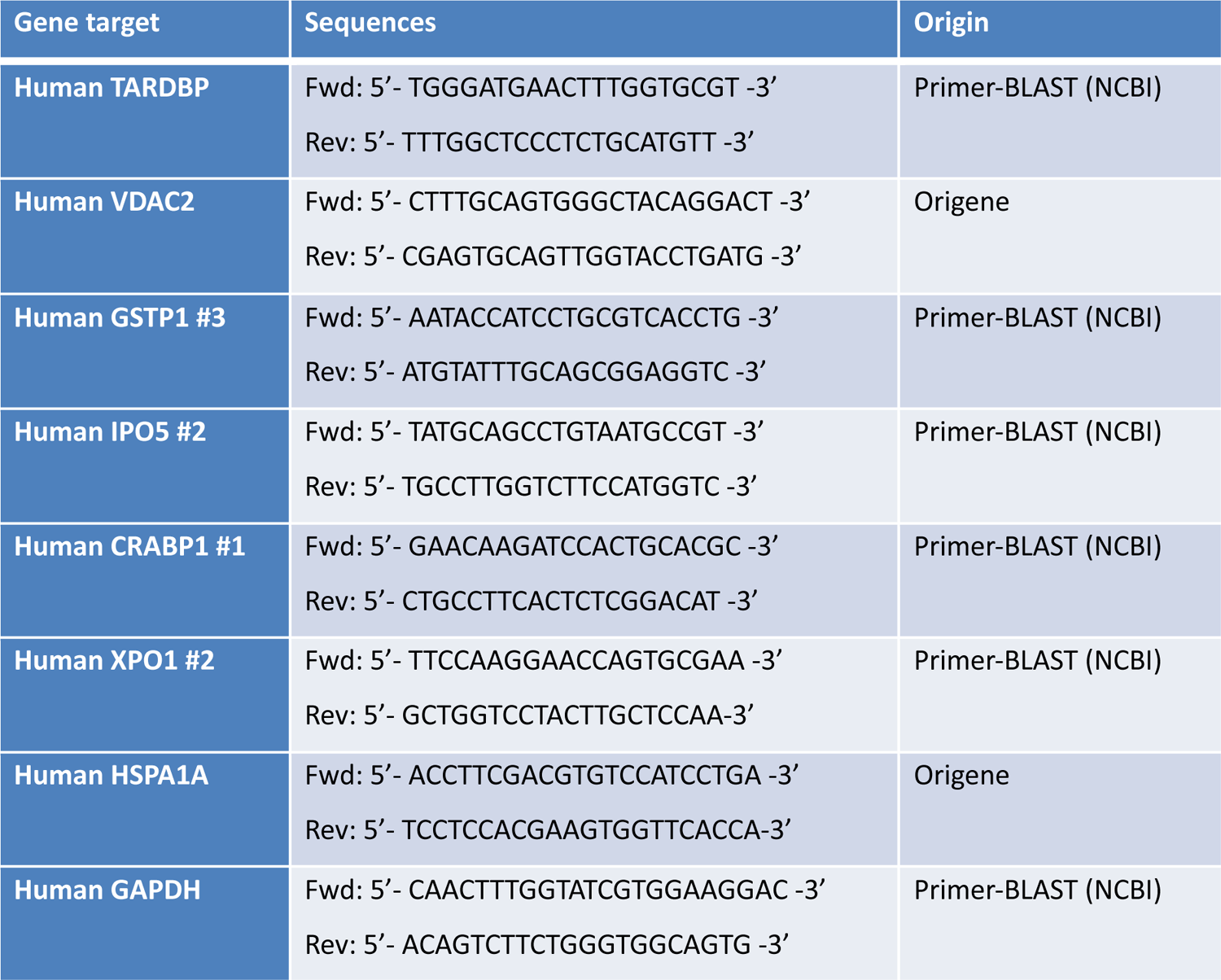
List of qPCR primers used in this study.

**Supplementary Table 2.** List of antibodies used in this study.

| Antibody | Host species | Dilution | Supplier |
| --- | --- | --- | --- |
| TDP-43 | Rabbit | 1:1,000 | Proteintech<br>#12892-1-AP |
| HSP-72 | Mouse | 1:1,000 | Merck<br>#386032 |
| eIF4A | Rabbit | 1:1,000 | Cell Signaling<br>#2490 |
| eEF2 | Rabbit | 1:10,000 | Abcam<br>Ab#75748 |
| RPL26 | Rabbit | 1:500 | Sigma<br>#R0655 |
| RPL29 | Rabbit | 1:2,000 | A gift from Prof. Stuart A. Wilson<br>(University of Sheffield) |
| GSTP1 | Rabbit | 1:2,000 | Proteintech<br>#15902-1-AP |
| VDAC 1-3 | Rabbit | 1:2,000 | Proteintech<br>#11663-1-AP |
| IPO5 | Rabbit | 1:2,000 | Invitrogen<br>#PA5-30076 |
| XPO1 | Rabbit | 1:2,000 | Abcam<br>#ab180144 |
| CRABP1 | Mouse | 1:1,000 | Abcam #ab2816 |
| α-Tubulin<br>(clone DM1A) | Mouse | 1:2,000 | Insight<br>#sc32293 |
| α-Tubulin<br>(11H10, for IF) | Rabbit | 1:2,000 | Cell Signaling<br>##2125 |
| GAPDH<br>(14C10) | Rabbit | 1:1,000 | Cell Signaling<br>mAb #2118 |

